# Bivalent COVID-19 vaccines boost the capacity of pre-existing SARS-CoV-2-specific memory B cells to cross-recognize Omicron subvariants

**DOI:** 10.1101/2024.03.20.585861

**Authors:** Holly A. Fryer, Daryl Geers, Lennert Gommers, Luca M. Zaeck, Ngoc H. Tan, Bernadette Jones-Freeman, Abraham Goorhuis, Douwe F. Postma, Leo G. Visser, P. Mark Hogarth, Marion P. G. Koopmans, Corine H. GeurtsvanKessel, Robyn E. O’Hehir, P. Hugo M. van der Kuy, Rory D. de Vries, Menno C. van Zelm

## Abstract

Bivalent COVID-19 vaccines comprising ancestral Wuhan-Hu-1 (WH1) and the Omicron BA.1 or BA.5 subvariant elicit enhanced serum antibody responses to emerging Omicron subvariants. We characterized the memory B-cell (Bmem) response following a fourth dose with a BA.1 or BA.5 bivalent vaccine, and compared the immunogenicity with a WH1 monovalent fourth dose. Healthcare workers previously immunized with mRNA or adenoviral vector monovalent vaccines were sampled before and one-month after a monovalent, BA.1 or BA.5 bivalent fourth dose COVID-19 vaccine. RBD-specific Bmem were quantified with an in-depth spectral flow cytometry panel including recombinant RBD proteins of the WH1, BA.1, BA.5, BQ.1.1, and XBB.1.5 variants. All recipients had slightly increased WH1 RBD-specific Bmem numbers. Recognition of Omicron subvariants was not enhanced following monovalent vaccination, while both bivalent vaccines significantly increased WH1 RBD-specific Bmem cross-recognition of all Omicron subvariants tested by flow cytometry. Thus, Omicron-based bivalent vaccines can improve recognition of descendent Omicron subvariants by pre-existing, WH1-specific Bmem, beyond that of a conventional, monovalent vaccine. This provides new insights into the capacity of variant-based mRNA booster vaccines to improve immune memory against emerging SARS-CoV-2 variants.

## INTRODUCTION

The mRNA- and adenoviral vector-based COVID-19 vaccines, encoding the Spike (S) protein of the ancestral Wuhan-Hu-1 lineage (WH1), are highly effective at preventing severe disease and hospitalization from SARS-CoV-2 (*1, 2*). However, the emergence of antigenically distinct Omicron subvariants in 2022 required the use of updated booster vaccinations to overcome reduced vaccine-induced neutralizing antibody (NAb) responses and maintain efficacy against emerging variants (*3–6*). Therefore, in addition to monovalent WH1 vaccines, fourth dose vaccinations were performed with bivalent vaccines that contain equal parts of mRNA encoding the WH1 and Omicron BA.1 or BA.5 S protein (*6–10*).

Bivalent vaccines proved significantly more effective at preventing infection and particularly severe disease or death from Omicron variants compared to monovalent WH1 vaccines (*11–13*). Bivalent boosters elicited equivalent levels of NAb against WH1 compared to monovalent vaccines, and increased NAb levels against the Omicron subvariant encoded by the bivalent vaccine, as well as descendant subvariants (*14–17*). Despite the induction of Omicron-specific immune responses, NAb levels against emerging subvariants are still significantly lower compared to WH1 (*14, 15, 18*). While NAb levels were initially considered a correlate of protection against COVID-19, the durable protection against severe disease is suggestive of a more prominent role for memory T- and B-cells. As the S receptor-binding domain (RBD) is the major target for NAb, quantification of RBD-specific memory B cells (Bmem) can be used as a correlate of long-term protection against severe COVID-19 (*2, 19–21*).

Circulating antigen-specific Bmem, detected in peripheral blood by flow cytometry, can be used to define the kinetics and phenotype of the S- and RBD-specific Bmem response to SARS-CoV-2 infection and vaccination (*19, 22–27*). Our group recently showed that a third monovalent mRNA dose boosted the frequency of WH1-specific Bmem binding Omicron BA.2 and BA.5 (*27*). Thus, the question remains whether the use of bivalent booster vaccines for the fourth dose enhances the recognition of Omicron subvariants compared to a monovalent WH1 vaccine, thereby broadening the SARS-CoV-2-specific immune responses. Here, we characterized and compared the NAb and Bmem responses following WH1 monovalent, BA.1 or BA.5 bivalent vaccination in a cohort of healthcare workers (HCW) from the Dutch SWITCH-ON study (*28*) and the Monash Immunology cohort.

## RESULTS

### Cohort characteristics

HCW were recruited from the Dutch SWITCH-ON study (*28*) and the Monash Immunology cohort (*25, 27*) for direct comparison of antibody and Bmem responses to monovalent, BA.1 bivalent, or BA.5 bivalent fourth dose COVID-19 vaccination (**Table 1**). Eighteen recipients of a monovalent booster (BNT162b2 or mRNA-1273), 33 BA.1 bivalent booster recipients (BNT162b2.BA1 or mRNA-1273.214), and 21 BA.5 bivalent booster recipients (BNT162b2.BA5 or mRNA-1273.222) were included (**Figure 1A**). The monovalent group comprised entirely Monash Immunology donors, while both bivalent groups were a combination of Monash and SWITCH-ON donors (**Table 1**, **Figure 1A**). Peripheral blood was sampled pre-dose four- and 28-days post-dose four (median 28 days, 27-49; **Table 1**, **Figure 1A**).

**Figure 1.**
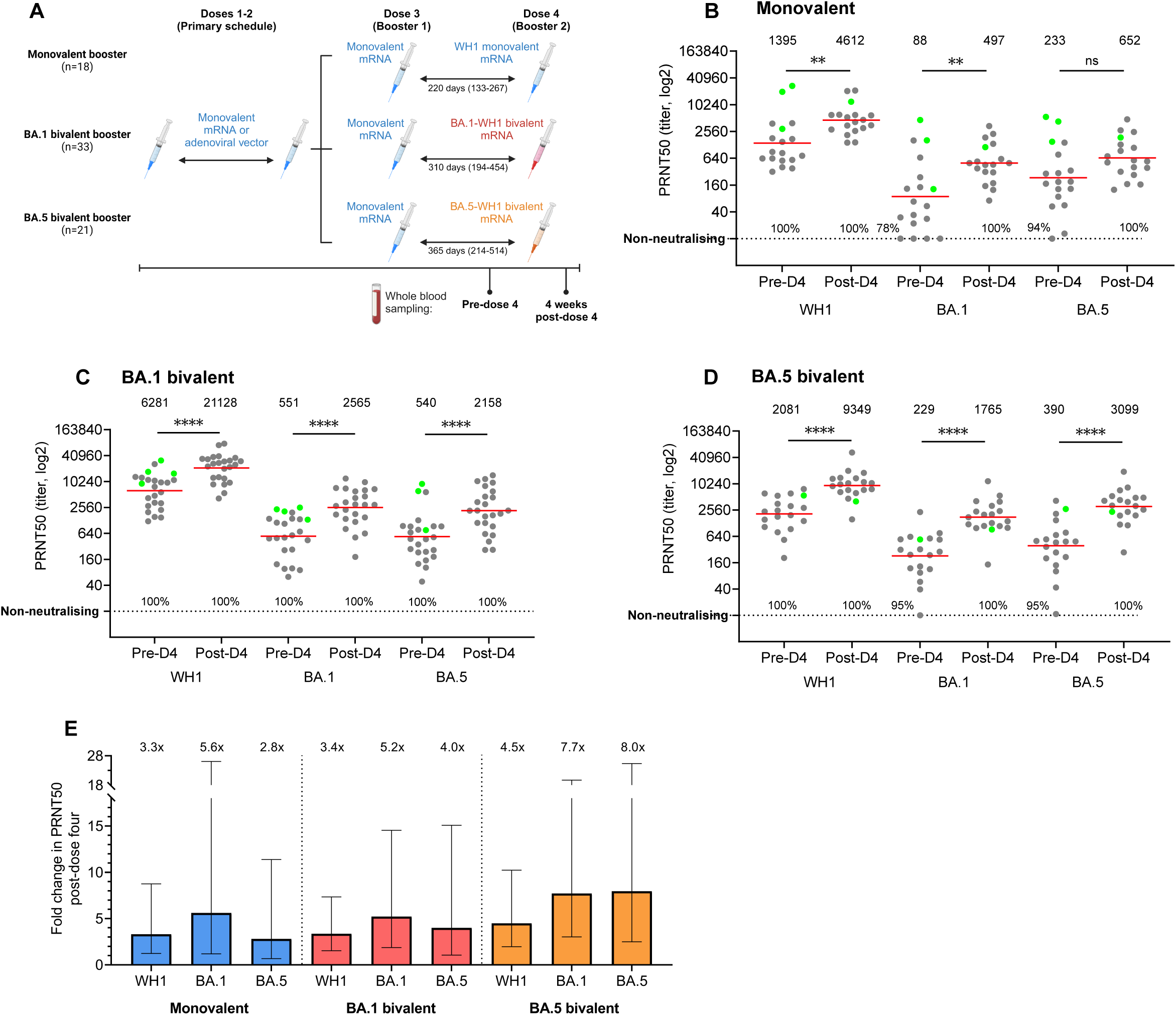
SARS-CoV-2 WH1, Omicron BA.1 and BA.5 neutralizing antibody responses elicited by monovalent, BA.1 bivalent, or BA.5 bivalent 4^th^ dose boosters. **(A)** Study design. Sampling was performed pre- and 4-weeks post-dose 4 (full cohort characteristics in **Table 1**). (B) PRNT50 NAb titers against WH1, BA.1, and BA.5 pre- and 4-weeks post-monovalent 4^th^ dose, (C) BA.1 bivalent 4^th^ dose, and (D) BA.5 bivalent 4^th^ dose. In (A-C), solid lines and values above panels indicate geometric means, horizontal dotted line denotes the neutralizing cutoff value of 10 for PRNT50, and percentages indicate the frequency of donors producing neutralizing antibody levels. (E) Fold changes in NAb titers against WH1, BA.1, and BA.5 4-weeks post-monovalent, BA.1 bivalent, or BA.5 bivalent 4^th^ dose. In (E), bars and values above panels indicate geometric means with geometric SDs. Monovalent: n=18; BA.1 bivalent: n=24, BA.5 bivalent: n=19. Green values indicate confirmed SARS-CoV-2 BTI prior to sampling. Wilcoxon matched-pairs signed rank test for paired data. **p<0.01, ****p<0.0001.

**Table 1.** Participant characteristics of the cohorts.

| Cohort detail | mRNA dose 4 group |  |  | Statistical comparisons |  |  |  |
| --- | --- | --- | --- | --- | --- | --- | --- |
|  | Monovalent WH1 | Bivalent BA.1 | Bivalent BA.5 | p-value |  |  |  |
|  | n=18 | n=33 | n=21 | Overall | Mono vs BA.1 | Mono vs BA.5 | BA.1 vs BA.5 |
| Recruitment center |  |  |  |  |  |  |  |
| Monash University | 100% (18/18) | 21% (7/33) | 9.5% (2/21) | <0.0001<sup>2</sup> |  |  |  |
| Erasmus MC | 0% (0/18) | 79% (26/33) | 90% (19/21) |  |  |  |  |
| Age (years; median with range) | 48.5 (32-65) | 46 (24-59) | 48 (22-65) | 0.6291<sup>1</sup> |  |  |  |
| Sex (%F) | 74 | 88 | 67 | 0.1519<sup>2</sup> |  |  |  |
| Vaccination characteristics |  |  |  |  |  |  |  |
| Primary vaccination/first 2 doses |  |  |  | 0.0232<sup>2</sup> | 0.016<sup>2</sup> | 0.0428<sup>2</sup> | >0.9999<sup>2</sup> |
| - mRNA | 3 (17%) | 15 (45%) | 11 (52%) |  |  |  |  |
| - ChAdOx1 | 15 (83%) | - | - |  |  |  |  |
| - Ad26.COV2.S* | - | 18 (55%) | 10 (48%) |  |  |  |  |
| 1<sup>st</sup> booster/3<sup>rd</sup> dose |  |  |  | - |  |  |  |
| - BNT162b2 | 16 (89%) | 33 (100%) | 21 (100%) |  |  |  |  |
| - ChAdOx1 | 2 (11%) | - | - |  |  |  |  |
| 2<sup>nd</sup> booster/4<sup>th</sup> dose |  |  |  | - |  |  |  |
| - BNT162b2 | 13 (72%) | - | - |  |  |  |  |
| - mRNA-1273 | 3 (17%) | - | - |  |  |  |  |
| - Novavax | 2 (11%) | - | - |  |  |  |  |
| - BNT162b2.BA1 | - | 9 (27%) | - |  |  |  |  |
| - mRNA-1273.214 | - | 24 (73%)<sup>4</sup> | - |  |  |  |  |
| - BNT162b2.BA5 | - | - | 11 (52%)<sup>5</sup> |  |  |  |  |
| - mRNA-1273.222 | - | - | 10 (48%)<sup>6</sup> |  |  |  |  |
| **Timing of dose 4** (days post-dose 3;<br>median with range) | 220 (133-267) | 310 (194-454) | 365 (214-533) | **<0.0001<sup>1</sup>** | **0.0001<sup>1</sup>** | **<0.0001<sup>1</sup>** | **0.0012<sup>1</sup>** |
| **Timing of sampling (days post-vaccination; median with range)** |  |  |  |  |  |  |  |
| **Pre-dose 4** (since dose 3) | 180.5 (129-191) | 308 (176-448) | 365 (214-514) | **<0.0001<sup>1</sup>** | **0.0001<sup>1</sup>** | **<0.0001<sup>1</sup>** | **0.0001<sup>1</sup>** |
| **Post dose 4** | 31.5 (28-48) | 28 (27-49) | 28 (27-30) | **<0.0001<sup>1</sup>** | **<0.0001<sup>1</sup>** | **<0.0001<sup>1</sup>** | **>0.9999<sup>1</sup>** |
| **Confirmed SARS-CoV-2 breakthrough infection (% infected, based on self-reporting and confirmed with NCP serology)** |  |  |  |  |  |  |  |
| **Any time before pre-dose 4 sampling** | 22% (4/18) | 70% (23/33) | 81% (17/21) | **0.0003<sup>2</sup>** |  |  |  |
| **Timing of BTI** (days before pre-dose 4<br>sampling; median with range) | 131.5 (61-148) | 238 (60-915) | 288 (169-938) | **0.0026<sup>1</sup>** | 0.0830<sup>1</sup> | **0.0028<sup>1</sup>** | 0.1319<sup>1</sup> |
| **Within 6 months of pre-dose 4**<br>**timepoint** | 17% (3/18) | 24% (8/33) | 5% (1/21) | 0.1732<sup>2**</sup> |  |  |  |
| **Between pre- and post-dose 4**<br>**timepoint** | 6% (1/18) | 6% (2/33) | 5% (1/21) | 0.9796<sup>2**</sup> |  |  |  |
| **Total B-cell count (absolute numbers in cells/μL; median with IQR)<sup>3</sup>** |  |  |  |  |  |  |  |
| **Pre-dose 4** | 203 (148-276) | 222 (159-288) | 138 (120-212) | **0.0057<sup>1</sup>** | >0.9999<sup>1</sup> | 0.0890<sup>1</sup> | **0.0047<sup>1</sup>** |
| **Post-dose 4** | 225 (183-253) | 189 (133-288) | 142 (121-181) | **0.0299<sup>1</sup>** | >0.9999<sup>1</sup> | **0.0425<sup>1</sup>** | 0.0946<sup>1</sup> |
| **Total Bmem count (absolute numbers in cells/μL; median with IQR)<sup>3</sup>** |  |  |  |  |  |  |  |
| **Pre-dose 4** | 90 (38-114) | 77 (43-101) | 47 (31-59) | **0.0337<sup>1</sup>** | >0.9999<sup>1</sup> | 0.0793<sup>1</sup> | 0.0590<sup>1</sup> |
| **Post-dose 4** | 78 (53-104) | 63 (40-106) | 42 (28-64) | **0.0191<sup>1</sup>** | 0.9133<sup>1</sup> | **0.0195<sup>1</sup>** | 0.1201<sup>1</sup> |

<sup>1</sup>Kruskal-Wallis test with Dunn's multiple comparisons; <sup>2</sup>Chi-squared test; <sup>3</sup>Data in **Supplementary Figure 3A, B**
\*Within BA.1 bivalent cohort, 5/18 Ad26.COV2.S recipients only received a single dose for their primary schedule, 1/18 received 2 doses, and 6 received 1 Ad26.COV.S dose followed by 1 mRNA vaccine dose. Within BA.5 bivalent cohort, 1/10 Ad26.COV2.S recipients only received a single dose for their primary schedule, 3/10 received 2 doses, and 6 received 1 Ad26.COV.S dose followed by 1 mRNA vaccine dose.
\*\*Low expected values for chi-squared test
Significant differences (p<0.05) in **bold**

There were no significant differences in the age and sex demographics between each of the three vaccine groups, with similar female preponderance. A significantly higher proportion of participants in the monovalent booster group had received primary vaccination (doses 1-2; 83%) with an adenoviral vector vaccine rather than with an mRNA vaccine as compared to both bivalent groups (48-55%). The time interval between the third and fourth vaccine doses was significantly different between the three groups, ranging from a median of 220 days (range 133-267) for the monovalent group, 310 days (194–454) for BA.1 bivalent and 365 (214–533) for BA.5 bivalent (**Table 1**). More donors in both bivalent groups had experienced a confirmed breakthrough infection (BTI) prior to dose four than the monovalent group, likely because the bivalent recipients had longer time intervals between third and fourth doses and thus more chance of infection (**Table 1**). The majority of these BTIs were reported between late 2021 and 2023, when Omicron subvariants were dominant (*4*). Whilst failing to reach significance, the median interval between BTIs and the pre-dose four sampling was slightly shorter in the BA.1 group than the BA.5 group, likely because the BA.5 group received their booster at a later timepoint due to the delayed introduction of the BA.5 bivalent vaccines (*29*).

### Increased neutralization of SARS-CoV-2 Omicron subvariants after a fourth dose booster

Plasma NAb titers against the SARS-CoV-2 WH1, Omicron BA.1, and BA.5 viruses were measured using a plaque reduction neutralization test (PRNT) before and four weeks after dose four. All donors, irrespective of vaccine type, had detectable NAb against WH1, BA.1, and BA.5 after dose four (**Figure 1B-D**). The monovalent vaccine elicited a significant increase in NAb titers against WH1 and BA.1 (**Figure 1B**), while the BA.1 and BA.5 bivalent vaccines elicited significant increases in NAb against WH1, BA.1 and BA.5. (**Figure 1C-D**). WH1 NAb titers were ∼3-4-fold higher after either monovalent or bivalent vaccination (**Figure 1E**). In contrast, the fold increases in BA.5 NAb titers were greater after the bivalent boosters than the monovalent booster, with the BA.5 bivalent booster eliciting the largest fold increases in all NAb titers (**Figure 1E**).

At baseline (pre-dose four), we observed higher NAb titers in the BA.1 bivalent cohort than both the monovalent and BA.5 bivalent cohorts. Due to the SWITCH-ON trial structure, the BA.1 bivalent cohort received their fourth dose booster three months earlier than the BA.5 bivalent cohort, which accounts for the higher titers (*30*). Individuals with a confirmed SARS-CoV-2 BTI within six months before a sampling timepoint tended to have higher NAb titers against WH1, BA.1, and BA.5, which may be contributing to the higher baseline and post-dose four NAb levels in the bivalent groups (**Figure 1B-D**). Overall, robust neutralization of WH1 as well as Omicron subvariants four weeks after a monovalent, BA.1 or BA.5 bivalent fourth dose booster was detected, with the bivalent BA.5 vaccine eliciting the greatest increases in NAb against Omicron BA.1 and BA.5.

### Bivalent vaccines boosted RBD-specific Bmem recognizing the vaccine Omicron subvariant

To evaluate the capacity of each vaccine to boost Bmem specific for the variants encoded by each vaccine, total, WH1, Omicron BA.1, and BA.5 RBD-specific Bmem were quantified and compared pre- and four-weeks post-dose four using flow cytometry (**Figure 2**; **Supplementary Figure 1)**. In the monovalent and BA.1 bivalent booster groups, B cells specific for WH1 and BA.1 RBDs were identified through double-discrimination to exclude any B cells binding to a fluorochrome. In the BA.5 bivalent booster group, double-discrimination was performed for WH1 and BA.5 RBDs (**Figure 2A**). Within RBD-specific B cells, mature Bmem were defined as CD38^dim^ and through subsequent exclusion of naive IgD^+^CD27^-^ B-cells (**Figure 2B**). The BA.1 bivalent group had more WH1 RBD-specific Bmem cells than the monovalent and BA.5 bivalent group, both at baseline (pre-dose four) and at four-weeks post-dose four (**Figure 2C**). This is potentially due to the higher frequency of recent BTIs, which may have impacted RBD-specific Bmem numbers (**Supplementary Figure 2**). Absolute numbers of WH1 RBD-specific Bmem significantly increased after both the BA.1 and BA.5 bivalent boosters (**Figure 2C**), but not after the monovalent booster. The median fold changes in WH1 RBD-specific Bmem numbers were similar between the three booster vaccine types (**Supplementary Table 1**), suggesting similar effects, which might not be significant in the monovalent group due to the smaller sample size.

**Figure 2.**
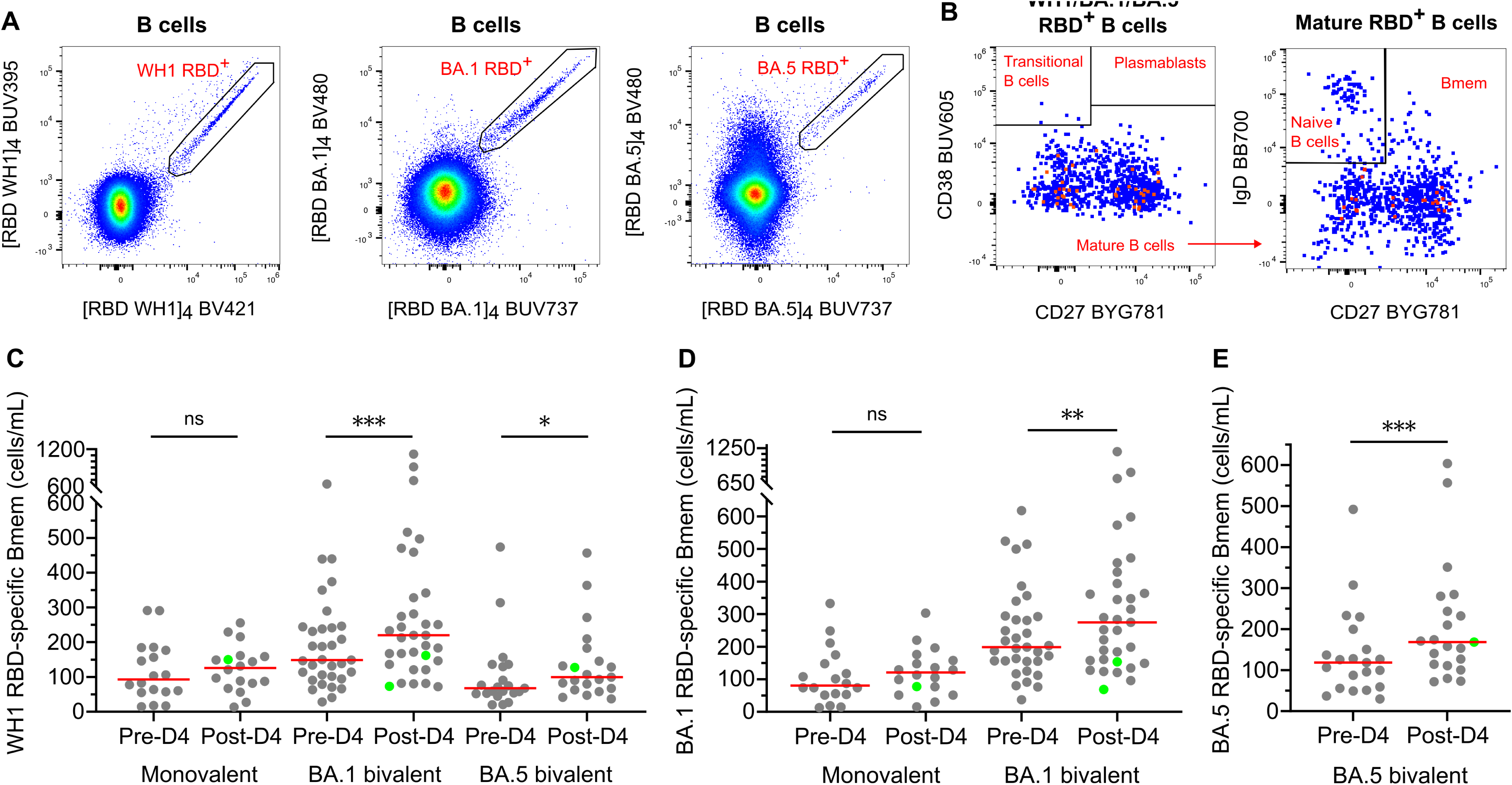
Significant increases in RBD-specific Bmem after a BA.1 or BA.5 bivalent 4^th^ dose booster. (**A**) Gating strategy for double-discrimination of WH1, BA.1, and BA.5 RBD-specific B cells by gating B cells double-positive for WH1 RBD, BA.1 RBD, or BA.5 RBD, respectively. (**B**) Sequential gating for mature B cells and memory B cells (Bmem) within RBD-specific B-cell populations. (**C**) Absolute numbers of WH1 RBD-specific Bmem pre- and 4-weeks post-monovalent, BA.1 bivalent, or BA.5 bivalent 4^th^ doses. (**D**) Absolute numbers of BA.1 RBD-specific Bmem pre- and 4-weeks post-monovalent or BA.1 bivalent 4^th^ doses. (**E**) Absolute numbers of BA.5 RBD-specific Bmem pre- and 4-weeks post-BA.5 bivalent 4^th^ dose. Monovalent dose 4, n=18; BA.1 bivalent dose 4, n=33; BA.5 bivalent dose 4, n=21. Solid lines depict medians. Green dots denote confirmed SARS-CoV-2 BTI between pre- and 4-weeks post-dose 4 sampling. Wilcoxon matched-pairs signed rank test for paired data. *p<0.05, **p<0.01, ***p<0.001.

The BA.1 and BA.5 bivalent boosters significantly increased the numbers of BA.1 or BA.5 RBD-specific Bmem, respectively (**Figure 2D, E**). There was no significant change in BA.1 RBD-specific Bmem after a monovalent booster, but the fold changes in median BA.1 RBD-specific Bmem after a monovalent and BA.1 bivalent booster were similar (**Supplementary Table 1**). The fold increase in BA.1 RBD-specific Bmem after a BA.1 bivalent booster was similar to the increase in BA.5 RBD-specific Bmem after a BA.5 bivalent booster (**Supplementary Table 1**).

### RBD-specific Bmem showed signs of recent activation after monovalent and bivalent boosters

The activation profile of RBD-specific Bmem was defined pre- and post-dose four through expression of cell-surface markers (**Figure 3; Supplemental Figure 3**). CD71 expression on Bmem is a marker of recent activation and proliferation, as is low CD21 expression due to downregulation upon antigen recognition (*31, 32*). Four weeks after a monovalent, BA.1 bivalent, or BA.5 bivalent booster, frequencies of CD71^+^CD38^dim^ WH1 RBD-specific Bmem increased significantly in the BA.1 and BA.5 bivalent booster recipients (**Figure 3A, B**). Frequencies of CD71^+^CD38^dim^ WH1 RBD-specific Bmem were not different between monovalent, BA.1 bivalent, and BA.5 bivalent booster recipients at either timepoint. Frequencies of CD21^lo^ WH1 RBD-specific Bmem increased after all fourth dose boosters, and there were no significant differences between the groups at either timepoint (**Figure 3C, D**).

**Figure 3.**
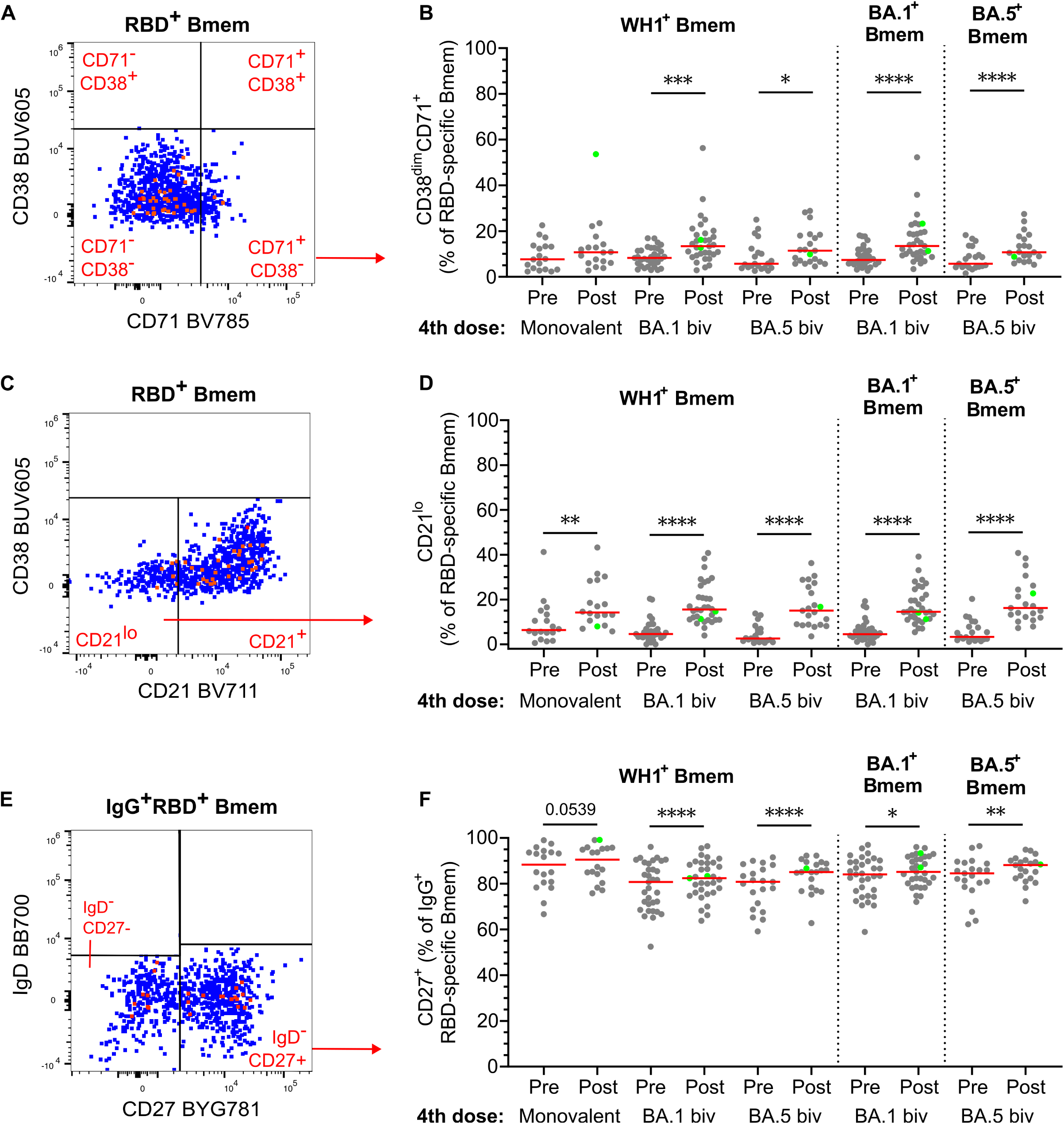
Activated RBD-specific Bmem following monovalent, BA.1 bivalent, and BA.5 bivalent 4^th^ dose vaccination. (A-B) CD38^dim^CD71^+^ events within RBD-specific Bmem. **(C-D)** CD21^lo^CD38^dim^ events within RBD-specific Bmem. **(E-F)** CD27^+^ events within IgG^+^ RBD-specific Bmem. Monovalent, n=18; BA.1 bivalent, n=33; BA.5 bivalent, n=21. Solid lines depict medians. Green dots denote confirmed SARS-CoV-2 BTI between pre- and 4-weeks post-dose 4 sampling. Wilcoxon matched-pairs signed rank test for paired data. Only significant differences shown. *p<0.05, **p<0.01, ***p<0.001, ****p<0.0001.

Within IgG^+^ WH1 RBD-specific Bmem, CD27 expression was measured as a marker of mature, germinal center (GC)-experienced, class-switched Bmem (**Figure 3E**) (*33*). The frequencies of CD27^+^IgG^+^ WH1 RBD-specific Bmem were higher after all three booster types, although not significant (p=0.054) for the monovalent group (**Figure 3F**). CD27^+^IgG^+^ WH1 RBD-specific Bmem frequencies before booster vaccination were significantly higher in the monovalent group than in both bivalent groups. The BA.1 and BA.5 bivalent boosters yielded similar increases in frequencies of CD38^dim^CD71^+^, CD21^lo^, and IgG^+^CD27^+^ BA.1 or BA.5 RBD-specific Bmem, respectively (**Figure 3B, D, F**).

Within the total Bmem population, no changes were observed in the frequencies of CD38^dim^CD71^+^, CD21^lo^, or IgG^+^CD27^+^ total Bmem (**Supplementary Figure 3C-E**). Thus, recipients of all three vaccines showed activation in their Bmem compartments with slightly higher increases following the bivalent boosters.

### mRNA-based priming had a sustained effect on IgG4^+^ Bmem after dose four

The Ig isotype and IgG subclass distributions of RBD-specific Bmem were evaluated pre- and four-weeks post-dose four booster (**Figure 4A**). At both timepoints in all booster type groups the majority of WH1 RBD-specific Bmem expressed IgG1 (70-88%; **Figure 4B**). The proportions of IgG1^+^ within WH1 RBD-specific Bmem were significantly higher than within total Bmem at both timepoints and in all groups (**Supplementary Figure 3F**). The monovalent and BA.1 bivalent boosters did not elicit any significant changes in Ig isotype distribution; however, the IgG3^+^ frequency tended to decrease in both groups, likely due to the slight expansion of the IgG1^+^ subset (**Figure 4B**). Within BA.1-specific Bmem following a BA.1 bivalent booster, the significant increase in IgG1^+^ BA.1-specific Bmem was accompanied by significant decreases in IgG3^+^, IgA^+^, and IgM^+^IgD^+^ subsets (**Figure 4B**). Following a BA.5 bivalent booster, the proportions of IgA^+^ Bmem within both WH1 and BA.5 RBD-specific Bmem were significantly lower, also likely due to the slight increase in IgG1^+^ frequencies (**Figure 4B**).

**Figure 4.**
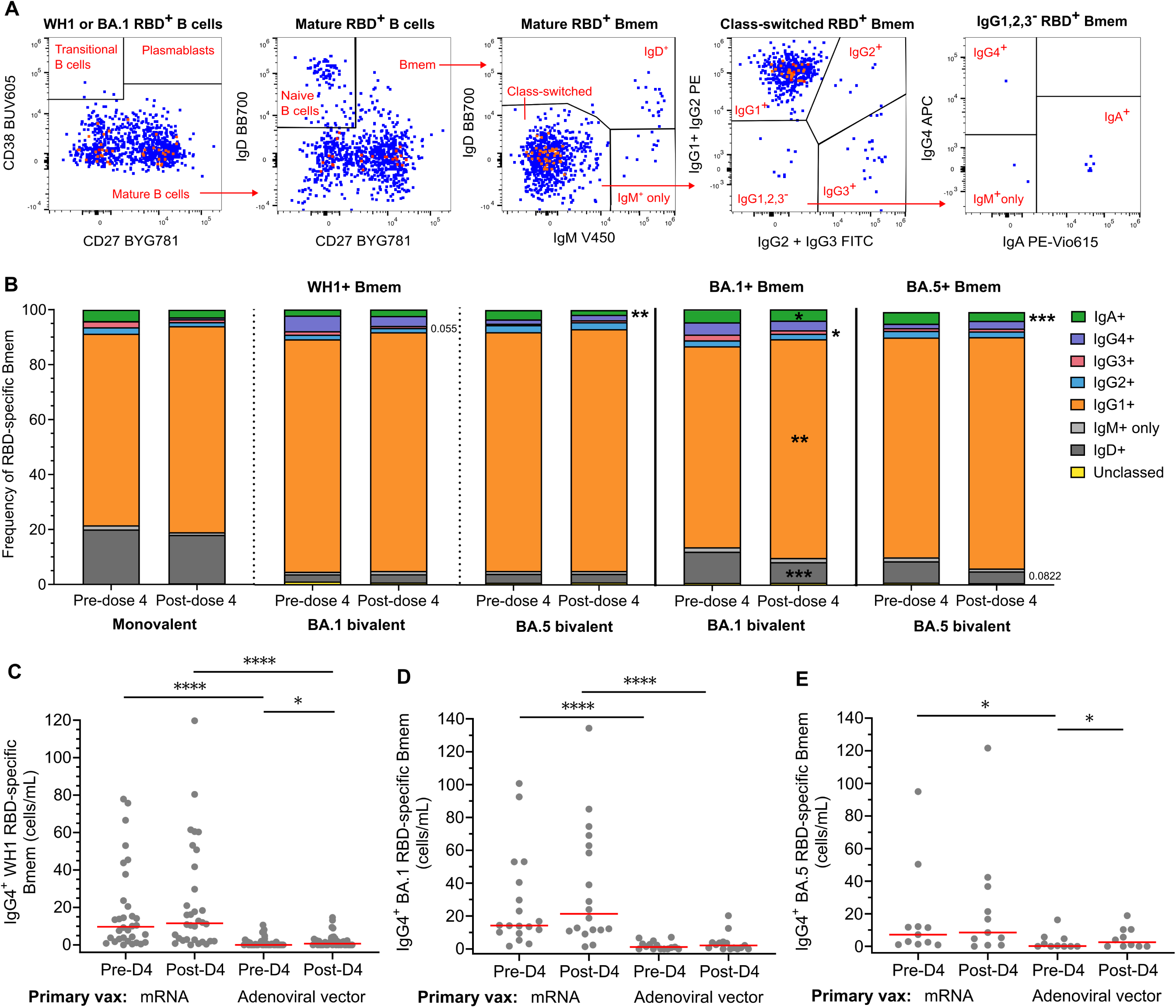
IgG subclass expression by RBD-specific Bmem after monovalent, BA.1 bivalent, or BA.5 bivalent 4^th^ doses. (**A**) Gating strategy of RBD-specific Bmem for Ig isotype and IgG subclass expression. (**B**) Distribution of Ig isotype and IgG subclass expressing subsets within WH1-specific Bmem, BA.1 RBD-specific Bmem, and BA.5 RBD-specific Bmem pre- and 4-weeks post-dose 4. Monovalent, n=18; BA.1 bivalent, n=33; BA.5 bivalent, n=21. (**C**) Absolute numbers of IgG4^+^ events within WH1, (**D**) BA.1, and (**E**) BA.5 RBD-specific Bmem pre- and post-dose 4 of all study subjects categorized based on priming with mRNA or adenoviral vector vaccines. (**C**) mRNA, n=32; adenoviral vector, n=40, (**D**) mRNA, n=18; adenoviral vector n=15, (**E**) mRNA, n=11; adenoviral vector, n=10. Solid lines depict medians. Mann-Whitney test for unpaired data and Wilcoxon matched-pairs signed rank test for paired data. Only significant differences shown. *p<0.05, ****p<0.0001.

We and others have recently reported that a third dose mRNA booster after double-dose mRNA priming elicits RBD-specific serum IgG4 and an IgG4^+^ Bmem population, which are both absent after mRNA boosting of an adenoviral vector-primed cohort (*27, 34, 35*). To evaluate if this effect is sustained after a fourth dose boost, we stratified all donors based on primary vaccination type (**Figure 4C**). mRNA-primed donors had significantly higher numbers of IgG4^+^ WH1 RBD-specific Bmem than adenoviral vector vaccine recipients before dose four (**Figure 4C**). These numbers were not affected by a fourth dose in mRNA recipients, but were significantly higher after dose four in adenoviral vector recipients, although still significantly lower than in mRNA-primed donors. Similar patterns were observed within BA.1 and BA.5 RBD-specific Bmem (**Figure 4D, E**).

### Bivalent vaccines broadened the recognition of Omicron subvariants by pre-existing WH1 RBD-specific Bmem

Next, we evaluated the capacity of WH1 RBD-specific Bmem to bind Omicron subvariants BA.1 and BA.5, as well as more recent sublineages BQ.1.1 (sublineage of BA.5) and XBB.1.5 (recombinant of two BA.2 subvariants) (*4, 36, 37*). The monovalent and BA.1 bivalent donors were evaluated for BA.1, BA.5, and BQ.1.1 binding within WH1-specific Bmem (**Figure 5A, Supplementary Table 2-Tube 2a**). For BA.5 bivalent donors, the panel was expanded to detect BA.1, BA.5, BQ.1.1, and XBB.1.5 binding within WH1-specific Bmem (**Figure 5B, Supplementary Table 2 - Tube 2b**). The numbers of WH1 RBD-specific Bmem that bound Omicron subvariant RBDs were significantly increased after both bivalent boosters, but not after a monovalent booster (**Figure 5C, D**).

**Figure 5.**
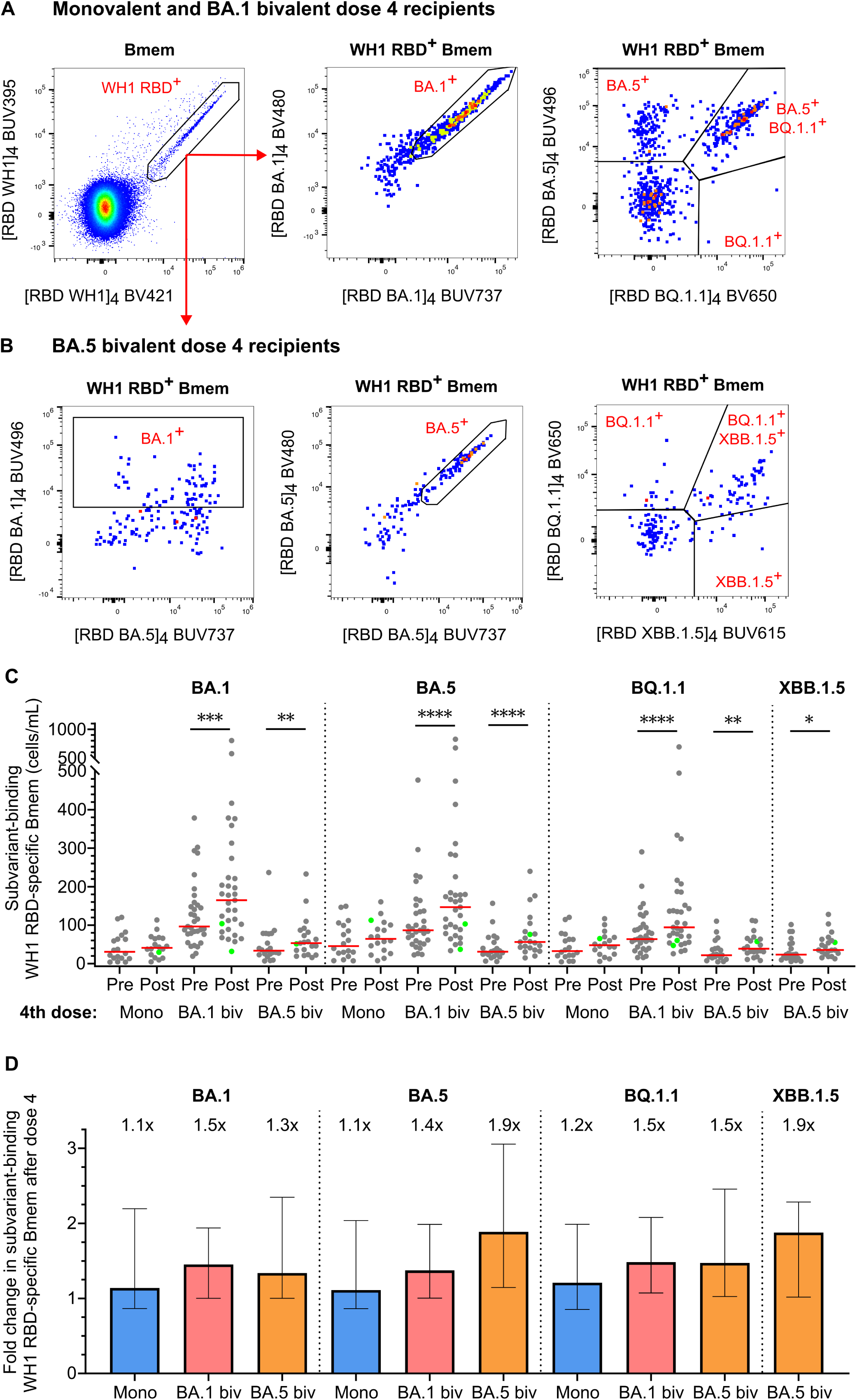
Enhanced capacity of WH1 RBD-specific Bmem to bind Omicron subvariants after boosting with bivalent vaccines. (**A**) Gating strategy to quantify WH1 RBD-specific Bmem that recognize Omicron BA.1, BA.5, and BA.1.1 in monovalent and BA.1 bivalent dose 4 recipients. Representative plots from a BA.1 bivalent booster recipient post-dose 4. (**B**) Gating strategy to quantify WH1 RBD-specific Bmem that recognize Omicron BA.1, BA.5, BA.1.1, and XBB.1.5 in BA.5 bivalent dose 4 recipients. Representative plots from a BA.5 bivalent booster recipient post-dose 4. (**C**) Absolute numbers of BA.1, BA.5, BQ.1.1, and XBB.1.5-specific cells within WH1 RBD-specific Bmem pre- and 4-weeks post-monovalent, BA.1 bivalent, or BA.5 bivalent dose 4. (**D**) Fold changes in WH1 RBD-specific Bmem binding Omicron BA.1, BA.5, BQ.1.1, or XBB.1.5 4-weeks post-monovalent, BA.1 bivalent, or BA.5 bivalent 4^th^ dose. In (**D**), bars and values above panels indicate medians with IQR. Monovalent, n=18; BA.1 bivalent, n=33; BA.5 bivalent, n=21. Solid lines indicate medians. Wilcoxon matched-pairs signed rank test for paired data. Only significant differences shown. *p<0.05, **p<0.01, ***p<0.001, ****p<0.0001.

As the BA.1 bivalent group was confounded by higher numbers of RBD-specific Bmem both pre- and post-dose four (**Figure 5C**), likely due to more recent Omicron BTIs (as discussed above), the fold increases were evaluated as well (**Figure 5D**). The BA.1 and BA.5 bivalent vaccines elicited similar fold increases for most variants, except for a larger increase in BA.5 binding for the BA.5 bivalent cohort. Thus, both the BA.1 and BA.5 bivalent vaccines elicited a greater capacity of WH1 RBD-specific Bmem to recognize Omicron subvariants, compared to the monovalent boosters.

### Omicron-only Bmem are increased by a BA.5 bivalent fourth dose booster

An early report indicated that boosting with a bivalent vaccine elicited Bmem with variant-only specificity, suggesting the recruitment of naive B cells with unique specificities into the booster response (*17*). We evaluated this for the BA.1 and BA.5 bivalent vaccines by detection of BA.1 and BA.5 specific Bmem, respectively, and then evaluation of the fraction that was negative for WH1 binding (**Figure 6A-C**). Pre-dose four, the BA.1 bivalent booster recipients had significantly higher numbers of BA.1-specific Bmem that did not bind WH1 compared to the monovalent group (**Figure 6D**). This was associated with the higher frequencies of BTIs during Omicron’s circulation prior to their fourth dose. The absolute number of BA.1^+^WH1^-^ Bmem did not change by four-weeks post-monovalent or BA.1 bivalent booster (**Figure 6D**). In contrast, the number of BA.5^+^WH1^-^ Bmem in was significantly increased following the fourth dose BA.5 bivalent booster (**Figure 6E**). Thus, in addition to expansion of WH1-specific Bmem with the capacity to bind BA.5, the BA.5 bivalent vaccine elicited expansion of BA.5-only binding Bmem.

**Figure 6.**
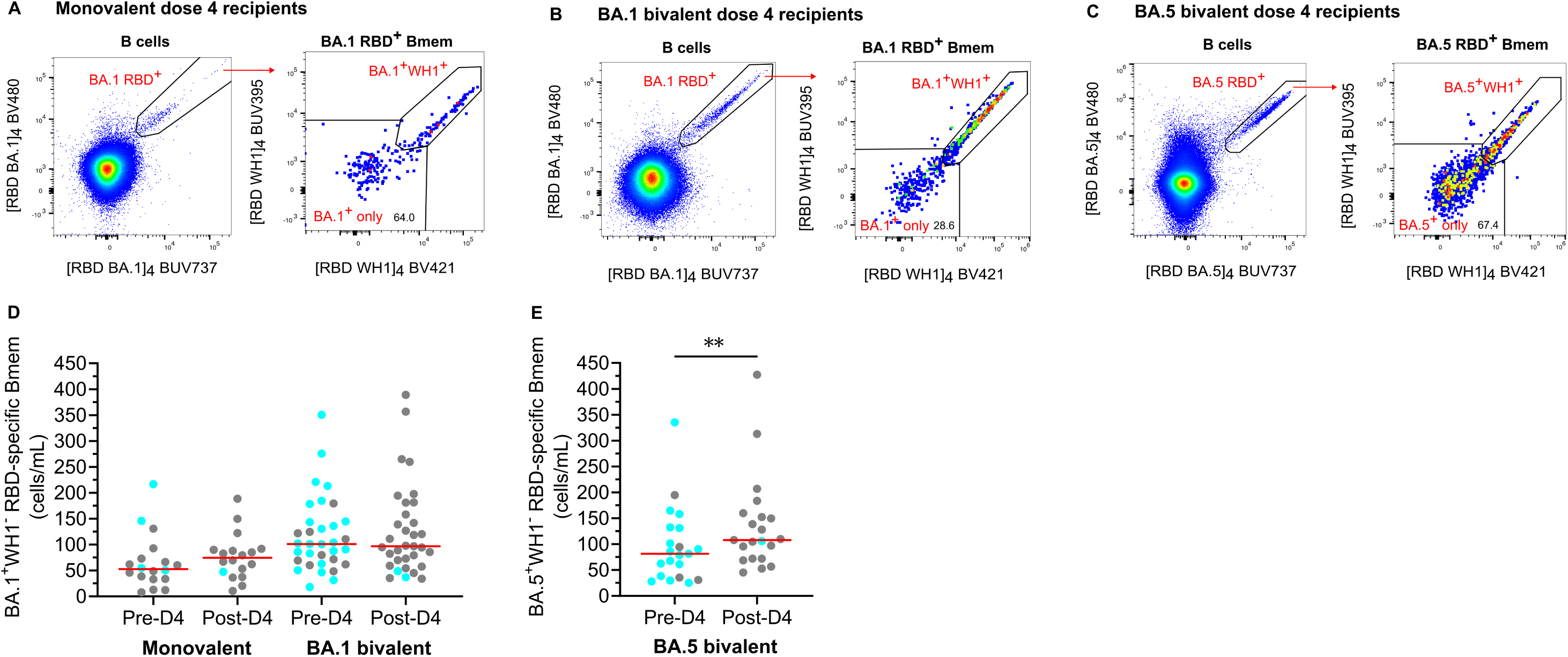
Omicron-only Bmem are increased by a BA.5 bivalent, but not a BA.1 bivalent or monovalent fourth dose booster. (**A-B**) Gating of BA.1^+^ only Bmem, negative for WH1 RBD binding, within BA.1 RBD-specific Bmem. Performed on monovalent and BA.1 bivalent dose 4 recipients. (**C**) Gating of BA.5+ only Bmem, negative for WH1 RBD binding, within BA.5 RBD-specific Bmem. Performed on BA.5 bivalent dose 4 recipients. (**D**) Absolute numbers of BA.1^+^WH1^-^ Bmem before and 4-weeks post-monovalent or BA.1 bivalent dose 4. (**E**) Absolute numbers of BA.5^+^WH1^-^ Bmem before and 4-weeks post-BA.5 bivalent dose 4. Monovalent, n=18; BA.1 bivalent, n=33; BA.5 bivalent, n=21. Blue dots denote any confirmed SARS-CoV-2 BTI before pre- or 4-weeks post-dose 4 sampling. Wilcoxon matched-pairs signed rank test for paired data. **p<0.01.

## DISCUSSION

We here performed for the first time, to our knowledge, a comparative evaluation of the capacity of monovalent WH1, bivalent BA.1 and bivalent BA.5 mRNA-based COVID-19 booster vaccinations to elicit Bmem responses that recognize emerging Omicron subvariants. Provided as fourth dose boosters, all three vaccine types boosted NAb levels against WH1, BA.1, and BA.5 variants. Monovalent and bivalent boosters similarly activated RBD-specific Bmem, and increased WH1 RBD-specific Bmem numbers. While recognition of Omicron subvariants was not increased in monovalent booster recipients, binding of BA.1, BA.5 and BQ.1.1 subvariants by WH1 RBD-specific Bmem was increased by both bivalent boosters, as was XBB.1.5 binding by the BA.5 bivalent booster. BA.5-only binding Bmem numbers were also boosted by the BA.5 vaccine booster, indicating its capacity to recruit new variant-only specific Bmem.

The WH1, BA.1, and BA.5 NAb titers of the bivalent vaccine recipients in our report displayed similar patterns as those from the SWITCH-ON trial (*30, 38*). We here extended these findings to report that both bivalent boosters elicited greater antibody responses than a monovalent booster, resulting in higher BA.1 and BA.5 NAb titers. This finding aligns with trials of the mRNA-1273.214 BA.1 bivalent vaccine, which was found to elicit superior NAb titers against its target antigen, Omicron BA.1, compared to the monovalent mRNA-1273 vaccine (*15, 16*). We also found that the BA.5 bivalent booster broadened variant NAb recognition the most, eliciting higher BA.5 NAb titers than the BA.1 bivalent and monovalent boosters, which confirms previous findings for this vaccine (*14, 15, 38–40*).

We found increases in the proportion of CD27^+^IgG^+^ RBD-specific Bmem at four-weeks after a monovalent or bivalent fourth dose, similar to trends we have shown following dose two and up to six-months post-dose three (*27, 41*). This indicates a continued maturation of Bmem to become resting over time after booster vaccination. We also report significant increases in the frequencies of CD21^lo^ and CD71^+^CD38^dim^ activated RBD-specific Bmem, illustrating the capacity of fourth dose boosters to re-activate a proportion of antigen-specific Bmem from quiescence. Others have observed a similar peak in CD21^lo^ S-specific B cells at four-weeks post-vaccination (*42*). It has been shown that CD21^lo^ Bmem have improved antigen-presenting capacity, which suggests that this CD21^lo^ RBD-specific Bmem population may be contributing to the vaccine response by activating T cells (*43*).

Our group previously reported a significantly larger proportion of IgG4^+^ Bmem in mRNA primary vaccine recipients compared to adenoviral vector recipients (*44*). Others have corroborated this expanded IgG4 response after two and three mRNA vaccine doses (*34, 35*). Notably, we now show a continued manifestation of this effect after an mRNA fourth dose, in the expression of IgG4 by WH1, BA.1, and BA.5 RBD-specific Bmem. One factor influencing this differential development of class-switching may be the difference in dosing interval, as mRNA-based primary vaccines were received three weeks apart compared to 12 weeks between ChAdOx1 adenoviral vector vaccines, and only a single dose was given to most Ad26.COV2.S recipients (*1, 45, 46*). Additionally, the mRNA-encoded S protein is stabilized by proline residues, while in adenoviral vector vaccines the DNA-encoded S protein can be truncated, and the S1 and S2 subunits are not stabilized and can be cleaved (*47*). As the S1 subunit contains the RBD, this difference in antigen structure may influence the development of the RBD-specific Bmem response (*17, 48*).

Overall, we found that the activation phenotypes and isotypes of RBD-specific Bmem were similar following either a monovalent or bivalent fourth dose booster. Therefore, the key difference in the Bmem response elicited by the bivalent boosters, compared to the conventional monovalent boosters, is the increase in breadth of variant binding. We found no significant increase in Bmem recognition of any Omicron subvariant RBD four weeks after a monovalent fourth dose booster. Our group previously observed that the frequency of Omicron BA.2 and BA.5 binding only increased by six-months post-dose three, so it is possible that measuring at the later timepoint is required to allow for Bmem maturation (*25, 27*). However, we found that four-weeks post-dose four the BA.1 and BA.5 bivalent vaccines boosted the ability of WH1 RBD-specific Bmem to bind antigenically distinct subvariants including those not contained in the bivalent vaccines, BQ.1.1 and XBB.1.5. This expands on previous analyses of NAb, which showed improved recognition of XBB and other related subvariants following the BA.5 bivalent mRNA vaccine or an Omicron BTI (*14, 15, 39, 49*). Therefore, we reveal novel evidence that cross-reactive Bmem binding both WH1 and Omicron subvariants are boosted by a bivalent fourth dose.

The enhanced ability of cross-reactive Bmem to bind Omicron after bivalent vaccination may be due to ongoing GC reactions that increase BCR affinity for variant RBDs. Exposure to viral variants through infection or vaccination is known to improve variant recognition by Bmem through continued maturation in the GC, linked to increased somatic hypermutations and higher cross-reactive BCR affinity (*50, 51*). There is evidence that this mechanism may be the cause of improved Bmem recognition of Omicron following booster vaccination, as bivalent vaccines have been shown to elicit prolonged GC B cell responses as well as BA.1- and BA.5-specific CD4^+^ T cells (*5, 17, 52–54*).

Neither the monovalent nor BA.1 bivalent boosters increased WH1-negative Omicron-specific Bmem, but BA.5-only Bmem did increase after a BA.5 bivalent booster. This is in line with the observed increase in neutralization breadth that was greatest following the BA.5 bivalent booster. There is previous evidence of a rare *de novo* Omicron-only binding Bmem population following Omicron-based monovalent vaccination; however, in the same study a majority of monoclonal antibodies isolated from Omicron S-specific Bmem were cross-reactive with the ancestral S protein (*17*). Therefore, the inclusion of the WH1 S protein in current bivalent vaccines may be limiting the development of these *de novo* populations, in a phenomenon known as immune imprinting or original antigenic sin. Pre-existing immune memory specific for the ancestral strain of a pathogen, elicited by primary vaccination or infection, can limit recruitment of naive B cells specific for variant epitopes through competition upon exposures with variant antigen (*11, 17, 25, 55*).

In May 2023, the WHO recommended the use of monovalent Omicron XBB vaccines in an effort to increase the breath of SARS-CoV-2 immunity (*4, 12, 17*). Phase 2/3 trials found that the XBB.1.5 monovalent mRNA vaccine elicited higher NAb titers against XBB.1.5 and XBB.1.16 than a bivalent XBB.1.5/BA.5 formulation (*17, 56*). Our current data show that the inclusion of Omicron vaccine antigens can enhance the breadth of Bmem binding to emergent subvariants, and exclusion of the WH1 antigen may reduce the limitations of immune imprinting (*17, 34*). Therefore, our findings support the use of monovalent variant-based mRNA vaccines going forward. However, preliminary data suggest that the recall of WH1-specific Bmem still dominates the response even after a monovalent XBB.1.5 booster (*57*).

There are limitations in the translational capacity of the study due to the predominance of females, the inclusion of only healthy adults under 65 years old, and the majority of donors being Caucasian. However, the study still provides baseline with which to compare the Bmem responses of high-risk populations including pediatric, elderly, immunodeficient, and immunocompromised individuals, which could help tailor their vaccine regimens and optimize their protection against emerging variants. Unavoidably, the monovalent cohort had a small sample size due to changes in Australian booster recommendations, resulting in a lower-powered group. Our inclusion of fold change analyses allowed us to detect some boosting effects of the monovalent fourth dose that may have not been otherwise significant.

Several factors may have contributed to the higher absolute numbers and frequencies of WH1 and Omicron RBD-specific Bmem in the BA.1 bivalent group. Firstly, the majority of the BA.1 bivalent group had at least one confirmed BTI (with Omicron) in the year prior to pre-dose four sampling, compared to only 22% of monovalent donors. These BTIs may have therefore elicited more Omicron-specific Bmem, including Omicron-only Bmem, resulting in the higher numbers at our baseline measures. Secondly, the BA.5 bivalent group received their fourth dose later than the BA.1 bivalent group, resulting in a slightly longer interval between their last BTI and pre-dose four sampling, which may have resulted in their lower Omicron-specific Bmem numbers.

Overall, Omicron BA.1 and BA.5 bivalent mRNA-based vaccines both increased the capacity of WH1 RBD-specific Bmem to bind all measured Omicron subvariants beyond that of a monovalent vaccine, showing that boosting with an antigenically distinct variant enhances the ability of pre-existing Bmem to bind to related subvariants. Our results reveal the cellular immune memory basis for understanding the higher degree of protection the bivalent boosters confer compared to monovalent WH1 COVID-19 vaccines, and supports the continued use of variant-based vaccines to prevent severe disease from emergent variants.

## MATERIALS AND METHODS

### Study design

From February 2021 to June 2023, healthy adults (18-65 years old, with no immunological or hematological disease) who received a monovalent, BA.1 bivalent, or BA.5 bivalent fourth dose COVID-19 booster were recruited to a research study conducted by Monash University at the Alfred Hospital (Australia) (**Table 1**). Additionally, HCW (18-65 years old) were recruited to the SWITCH-ON study, a multicenter randomized controlled trial involving four academic hospitals in the Netherlands and randomized to groups who received a fourth dose BA.1 bivalent COVID-19 booster in October 2022, or a BA.5 bivalent booster in December 2022, respectively. Full details can be found in the trial protocol (*29*). A combined total of 72 donors, 27 participants from the Monash University project and 45 participants from the SWITCH-ON study, were analyzed in this manuscript (**Table 1**). Following written informed consent, peripheral blood samples were collected pre-dose four booster, and four-weeks post-dose four booster. Blood samples were processed, as previously described, to perform TruCount analysis, and to isolate plasma or serum and PBMC for detailed immunological analysis (see below) (*24*). Demographic information including age, sex, prior vaccination dates and types, and SARS-CoV-2 infection status were collected throughout the studies. Reported SARS-CoV-2 breakthrough infections (BTIs) were confirmed with nucleocapsid protein (NCP)-specific IgG assays, as described previously (*24, 25, 27, 58*). The studies were conducted according to the Declaration of Helsinki and approved by local human research ethics committees (Monash Immunology cohort: Alfred Health ethics no. 32/21, Monash University project no. 72794; SWITCH-ON trial: Erasmus Medical Center Medical Ethics Review Committee, protocol no. MEC-2022-0462, and local review boards of participating centers, and registered at ClinicalTrials.gov, no. NCT05471440).

### PRNT assay

NAb were measured for all donor plasma samples using a plaque reduction neutralization test (PRNT), as described previously (*5, 30, 59*). Viruses were isolated and cultured from clinical specimens from the Department of Viroscience, Erasmus MC, and confirmed by next-generation sequencing: D614G (ancestral; GISAID: hCov-19/Netherlands/ZH-EMC-2498), Omicron BA.1 (GISAID: hCoV-19/Netherlands/LI-SQD-01032/2022), and Omicron BA.5 (EVAg: 010V-04723; hCovN19/Netherlands/ZHNEMCN5892) (*38*). Briefly, heat-inactivated serum was serially diluted two-fold in OptiMEM without FBS (Gibco). Four hundred PFU of each SARS-CoV-2 variant in an equal volume of OptiMEM were added to the diluted sera and incubated at 37°C for 1 hour. The serum-virus mixture was transferred to human airway Calu-3 cells (ATCC HTB-55) and incubated at 37°C for 8 hours. The cells were then fixed in 10% neutral-buffered formalin, permeabilized in 70% ethanol, and plaques stained with a polyclonal rabbit anti-SARS-CoV-2 nucleocapsid antibody (Sino Biological) and a secondary peroxidase-labelled goat-anti rabbit IgG antibody (Dako). The signals were developed with a precipitate-forming TMB substrate (TrueBlue, SeraCare/KPL) and the number of plaques per well was quantified with an ImmunoSpot Image Analyzer (CTL Europe GmbH). The 50% reduction titer (PRNT50) was estimated by calculating the proportionate distance between two dilutions from which the endpoint titer was calculated. An infection control (without serum) and positive serum control (Nanogam® 100 mg/mL, Sanquin) were included on every assay plate. When no neutralization was detected, the sample was assigned an arbitrary PRNT50 value of 10.

### Protein production

DNA constructs encoding the SARS-CoV-2 RBD of WH1, Omicron BA.1, BA.5, BQ.1.1, and XBB.1.5 were designed incorporating an N-terminal Fel d 1 leader sequence, a C-terminal AviTag for biotin ligase (BirA)-catalyzed biotinylation, and a 6-His tag for cobalt affinity column purification (*24, 25, 27*). The DNA construct encoding the SARS-CoV-2 WH1 NCP protein was generated with an N-terminal human Ig leader sequence and the same C-terminal AviTag and 6-His tag (*24*). The DNA constructs were cloned into a pCR3 plasmid and produced using the Expi293 Expression system (Thermo Fisher, Waltham, MA), then purified, biotinylated, and tetramerized, as described previously (*24, 25, 27*). This generated fluorescent tetramers [RBD WH1]_4_-BUV395, [RBD WH1]_4_-BV421 and [RBD BQ.1.1]_4_-BV650 which were used in both panel variations, as well as [RBD BA.1]_4_-BV480, [RBD BA.1]_4_-BUV737, [RBD BA.5]_4_-BUV496 for the panel used to analyze monovalent and BA.1 bivalent booster recipients, and [RBD BA.5]_4_-BV480, [RBD BA.5]_4_-BUV737, [RBD BA.1]_4_-BUV496, and [RBD XBB.1.5]_4_-BUV615 for the panel used to analyze BA.5 bivalent booster recipients (**Supplementary Tables 2 and 3**).

### Flow cytometry

#### Trucount

Absolute numbers of major leukocyte populations were determined for each peripheral blood sample as previously described (*24, 25, 60*). Briefly, 50µL of fresh whole blood was added to a BD Trucount tube (BD Biosciences, San Jose, CA, USA) and incubated with 20µL of the Multitest^TM^ 6-color TBNK reagent (BD Biosciences) containing CD3, CD4, CD8, CD19, CD16, CD45 and CD56 antibodies (**Supplementary Tables 2 and 3**) for 15 minutes at room temperature in the dark. Subsequently, cells were incubated with 1X BD Lysis Solution (BD Biosciences) for 15 minutes to lyse red blood cells. Samples were acquired on the BD FACSLyric analyzer and data were analyzed using FlowJo^TM^ Software v10.9.0 (BD Biosciences) as previously described (*24, 60*). Trucount data were then used to calculate the absolute numbers of RBD-specific Bmem subsets (*60*).

#### RBD-specific Bmem analysis

Fluorescent tetramers of WH1, Omicron BA.1, BA.5, and BQ.1.1 RBDs were incorporated into a 19-colour spectral flow cytometry panel to characterize the RBD-specific Bmem response elicited by a fourth dose booster in the monovalent and BA.1 bivalent fourth dose groups (**Supplementary Tables 2 and 3**). Due to the emergence of subsequent Omicron subvariants including XBB.1.5 by the time the BA.5 bivalent vaccine was distributed, the previous panel was modified for analysis of samples from BA.5 bivalent fourth dose recipients to include WH1, BA.1, BA.5, BQ.1.1, and XBB.1.5 RBD tetramers in a 20-colour panel (**Supplementary Table 2 and 3**). For each pre- and four-weeks post dose four sample, 10-15×10^6^ thawed PBMC were incubated at room temperature in the dark for 15 minutes in a total volume of 250µL with FACS buffer (PBS with 0.1% sodium azide and 0.2% BSA), fixable ViaDye Red, antibodies against surface markers and 5µg/mL each of each RBD tetramer (**Supplementary Tables 2 and 3**). In a separate tube, 1-5×10^6^ PBMCs were incubated at room temperature in the dark for 15 minutes in a total volume of 100µL with FACS buffer, fixable ViaDye Red, antibodies against surface markers and fluorochrome-conjugated streptavidin controls (**Supplementary Tables 2 and 3**). Cells were then washed with FACS buffer, fixed with 2% PFA for 20 minutes at room temperature in the dark, washed once more and acquired on the Cytek Aurora (Cytek Biosciences) using SpectroFlo® software v3.1. Data analysis was performed using FlowJo^TM^ Software v10.9.0 (gating strategy in **Supplementary Figure 1**).

### Statistical analysis

Absolute numbers of RBD-specific Bmem were calculated relative to the B cell counts measured by the Trucount protocol. GraphPad Prism (v9.5.1) software was used for statistical analyses. Unpaired data were analyzed using the Mann-Whitney test, paired data with the Wilcoxon signed-ranks test, data across multiple groups with the Kruskal-Wallis test with Dunn’s multiple comparisons, and categorical data with the Chi-squared test. p<0.05 was considered significant for all statistical tests.

## ACKNOWLEDGEMENTS

We thank the ARAFlowCore staff for training and assistance with flow cytometry, Ms Sandra Esparon, Ms Reema Bajaj, and Dr Bruce D Wines (Burnet Institute) for assistance with protein production, Ms Pei Mun Aui, Ms Ebony Blight, Mr Jack Edwards, Dr Gemma Hartley, Ms Shir Sun, Ms Alina Wang (Monash University), and Ms Susanne Bogers (Erasmus MC) for sample collection and preparation, and Ms Laura van Dijk (Erasmus MC) for PRNT assays.

## AUTHORS’ CONTRIBUTIONS

Study design: HAF, AG, DFP, LGV, PMH, REOH, CHGvK, PHMvdK, RDdV and MCvZ; Performed experiments: HAF, DG, LG, LMZ, NHT, and MCvZ; Formal analysis: HAF, DG and LMZ; Subject recruitment/inclusion, vaccination, and sampling: NHT, BJF, AG, DFP, LGV and PHMvdK; Supervised the work: MPGK, CHGvK, PHMvdK, RDdV, MCvZ; Wrote the manuscript: HAF and MCvZ. All authors edited and approved the final version of the manuscript.

## DATA AND MATERIAL AVAILABILITY

Data and/or materials will be made available from the corresponding author upon reasonable request.

## ETHICS APPROVAL

This study was conducted according to the Declaration of Helsinki and approved by local human research ethics committees. Monash Immunology cohort: Alfred Health ethics no. 32/21, Monash University project no. 72794. The SWITCH ON trial study protocol was approved by the Erasmus Medical Center Medical Ethics Review Committee (protocol no. MEC-2022-0462), and local review boards of participating centers, and was registered at ClinicalTrials.gov (NCT05471440).

## CONSENT TO PARTICIPATE

Written informed consent was obtained from all individual participants prior to inclusion in the study.

## REFERENCES

1. F. P. Polack, S. J. Thomas, N. Kitchin, J. Absalon, A. Gurtman, S. Lockhart, J. L. Perez, G. Pérez Marc, E. D. Moreira, C. Zerbini, R. Bailey, K. A. Swanson, S. Roychoudhury, K. Koury, P. Li, W. V. Kalina, D. Cooper, R. W. Frenck, L. L. Hammitt, Ö. Türeci, H. Nell, A. Schaefer, S. Ünal, D. B. Tresnan, S. Mather, P. R. Dormitzer, U. Şahin, K. U. Jansen, and W. C. Gruber, Safety and Efficacy of the BNT162b2 mRNA Covid-19 Vaccine. New England Journal of Medicine 383, 2603–2615 (2020).

2. P. M. Folegatti, K. J. Ewer, P. K. Aley, B. Angus, S. Becker, S. Belij-Rammerstorfer, D. Bellamy, S. Bibi, M. Bittaye, E. A. Clutterbuck, C. Dold, S. N. Faust, A. Finn, A. L. Flaxman, B. Hallis, P. Heath, D. Jenkin, R. Lazarus, R. Makinson, A. M. Minassian, K. M. Pollock, M. Ramasamy, H. Robinson, M. Snape, R. Tarrant, M. Voysey, C. Green, A. D. Douglas, A. V. S. Hill, T. Lambe, S. C. Gilbert, and A. J. Pollard, Safety and immunogenicity of the ChAdOx1 nCoV-19 vaccine against SARS-CoV-2: a preliminary report of a phase 1/2, single-blind, randomised controlled trial. Lancet 396, 467–478 (2020).

3. N. Andrews, J. Stowe, F. Kirsebom, S. Toffa, T. Rickeard, E. Gallagher, C. Gower, M. Kall, N. Groves, A.-M. O’Connell, D. Simons, P. B. Blomquist, A. Zaidi, S. Nash, N. Iwani Binti Abdul Aziz, S. Thelwall, G. Dabrera, R. Myers, G. Amirthalingam, S. Gharbia, J. C. Barrett, R. Elson, S. N. Ladhani, N. Ferguson, M. Zambon, C. N. J. Campbell, K. Brown, S. Hopkins, M. Chand, M. Ramsay, and J. Lopez Bernal, Covid-19 Vaccine Effectiveness against the Omicron (B.1.1.529) Variant. New England Journal of Medicine 386, 1532–1546 (2022).

4. W. H. Organisation. 2023. Statement on the antigen composition of COVID-19 vaccines, Vol. 2023.

5. T. Arashiro, Y. Arima, J. Kuramochi, H. Muraoka, A. Sato, K. Chubachi, K. Oba, A. Yanai, H. Arioka, Y. Uehara, G. Ihara, Y. Kato, N. Yanagisawa, Y. Nagura, H. Yanai, A. Ueda, A. Numata, H. Kato, H. Oka, Y. Nishida, K. Ishii, T. Ooki, Y. Nidaira, T. Asami, T. Jinta, A. Nakamura, D. Taniyama, K. Yamamoto, K. Tanaka, K. Ueshima, T. Fuwa, A. Stucky, T. Suzuki, C. Smith, M. Hibberd, K. Ariyoshi, and M. Suzuki, Immune escape and waning immunity of COVID-19 monovalent mRNA vaccines against symptomatic infection with BA.1/BA.2 and BA.5 in Japan. Vaccine 41, 6969–6979 (2023).

6. Australian Government Department of Health. 2022. ATAGI recommendations on the use of a booster dose of COVID-19 vaccine, Vol. 2022.

7. Australian Government Department of Health. 2023. ATAGI recommendations on use of the Pfizer bivalent (Original/Omicron BA.4/5) COVID-19 vaccine, Vol. 2023.

8. European Medicines Agency. 2022. ECDC-EMA statement on booster vaccination with Omicron adapted bivalent COVID-19 vaccines.

9. Australian Government Department of Health. 2022. ATAGI recommendations on use of the Pfizer bivalent (Original/Omicron BA.1) COVID-19 vaccine, Vol. 2023.

10. S. Chalkias, C. Harper, K. Vrbicky, S. R. Walsh, B. Essink, A. Brosz, N. McGhee, J. E. Tomassini, X. Chen, Y. Chang, A. Sutherland, D. C. Montefiori, B. Girard, D. K. Edwards, J. Feng, H. Zhou, L. R. Baden, J. M. Miller, and R. Das, A Bivalent Omicron-Containing Booster Vaccine against Covid-19. New England Journal of Medicine, (2022).

11. M. Aguilar-Bretones, R. A. M. Fouchier, M. P. G. Koopmans, and G. P. van Nierop, Impact of antigenic evolution and original antigenic sin on SARS-CoV-2 immunity. The Journal of Clinical Investigation 133, (2023).

12. R. Arbel, A. Peretz, R. Sergienko, M. Friger, T. Beckenstein, H. Duskin-Bitan, S. Yaron, A. Hammerman, N. Bilenko, and D. Netzer, Effectiveness of a bivalent mRNA vaccine booster dose to prevent severe COVID-19 outcomes: a retrospective cohort study. Lancet Infect Dis 23, 914–921 (2023).

13. N. W. Andersson, E. M. Thiesson, U. Baum, N. Pihlström, J. Starrfelt, K. Faksová, E. Poukka, H. Meijerink, R. Ljung, and A. Hviid, Comparative effectiveness of bivalent BA.4-5 and BA.1 mRNA booster vaccines among adults aged ≥50 years in Nordic countries: nationwide cohort study. BMJ 382, e075286 (2023).

14. C. Kurhade, J. Zou, H. Xia, M. Liu, H. C. Chang, P. Ren, X. Xie, and P.-Y. Shi, Low neutralization of SARS-CoV-2 Omicron BA.2.75.2, BQ.1.1, and XBB.1 by 4 doses of parental mRNA vaccine or a BA.5-bivalent booster. *bioRxiv*, 2022.2010.2031.514580 (2022).

15. S. Chalkias, N. McGhee, J. L. Whatley, B. Essink, A. Brosz, J. E. Tomassini, B. Girard, K. Wu, D. K. Edwards, A. Nasir, D. Lee, L. E. Avena, J. Feng, W. Deng, D. C. Montefiori, L. R. Baden, J. M. Miller, and R. Das, Safety and Immunogenicity of XBB.1.5-Containing mRNA Vaccines. *medRxiv*, 2023.2008.2022.23293434 (2023).

16. S. Chalkias, C. Harper, K. Vrbicky, S. R. Walsh, B. Essink, A. Brosz, N. McGhee, J. E. Tomassini, X. Chen, Y. Chang, A. Sutherland, D. C. Montefiori, B. Girard, D. K. Edwards, J. Feng, H. Zhou, L. R. Baden, J. M. Miller, and R. Das, A Bivalent Omicron-Containing Booster Vaccine against Covid-19. New England Journal of Medicine 387, 1279–1291 (2022).

17. W. B. Alsoussi, S. K. Malladi, J. Q. Zhou, Z. Liu, B. Ying, W. Kim, A. J. Schmitz, T. Lei, S. C. Horvath, A. J. Sturtz, K. M. McIntire, B. Evavold, F. Han, S. M. Scheaffer, I. F. Fox, S. F. Mirza, L. Parra-Rodriguez, R. Nachbagauer, B. Nestorova, S. Chalkias, C. W. Farnsworth, M. K. Klebert, I. Pusic, B. S. Strnad, W. D. Middleton, S. A. Teefey, S. P. J. Whelan, M. S. Diamond, R. Paris, J. A. O’Halloran, R. M. Presti, J. S. Turner, and A. H. Ellebedy, SARS-CoV-2 Omicron boosting induces de novo B cell response in humans. Nature 617, 592–598 (2023).

18. J. N. Faraone, P. Qu, N. Goodarzi, Y.-M. Zheng, C. Carlin, L. J. Saif, E. M. Oltz, K. Xu, D. Jones, R. J. Gumina, and S.-L. Liu, Immune evasion and membrane fusion of SARS-CoV-2 XBB subvariants EG.5.1 and XBB.2.3. Emerging Microbes & Infections 12, 2270069 (2023).

19. A. Sokal, G. Barba-Spaeth, I. Fernández, M. Broketa, I. Azzaoui, A. de La Selle, A. Vandenberghe, S. Fourati, A. Roeser, A. Meola, M. Bouvier-Alias, E. Crickx, L. Languille, M. Michel, B. Godeau, S. Gallien, G. Melica, Y. Nguyen, V. Zarrouk, F. Canoui-Poitrine, F. Pirenne, J. Mégret, J.-M. Pawlotsky, S. Fillatreau, P. Bruhns, F. A. Rey, J.-C. Weill, C.-A. Reynaud, P. Chappert, and M. Mahévas, mRNA vaccination of naive and COVID-19-recovered individuals elicits potent memory B cells that recognize SARS-CoV-2 variants. Immunity 54, 2893–2907 (2021).

20. A. K. Wheatley, J. A. Juno, J. J. Wang, K. J. Selva, A. Reynaldi, H.-X. Tan, W. S. Lee, K. M. Wragg, H. G. Kelly, R. Esterbauer, S. K. Davis, H. E. Kent, F. L. Mordant, T. E. Schlub, D. L. Gordon, D. S. Khoury, K. Subbarao, D. Cromer, T. P. Gordon, A. W. Chung, M. P. Davenport, and S. J. Kent, Evolution of immune responses to SARS-CoV-2 in mild-moderate COVID-19. Nature Communications 12, 1162 (2021).

21. N. Sherina, A. Piralla, L. Du, H. Wan, M. Kumagai-Braesch, J. Andréll, S. Braesch-Andersen, Cassaniti, E. Percivalle, A. Sarasini, F. Bergami, R. Di Martino, M. Colaneri, M. Vecchia, M. Sambo, V. Zuccaro, R. Bruno, M. Sachs, T. Oggionni, F. Meloni, H. Abolhassani, F. Bertoglio, M. Schubert, M. Byrne-Steele, J. Han, M. Hust, Y. Xue, L. Hammarström, F. Baldanti, H. Marcotte, and Q. Pan-Hammarström, Persistence of SARS-CoV-2-specific B and T cell responses in convalescent COVID-19 patients 6 months after the infection. Med 2, 281–295 (2021).

22. R. R. Goel, S. A. Apostolidis, M. M. Painter, D. Mathew, A. Pattekar, O. Kuthuru, S. Gouma, P. Hicks, W. Meng, A. M. Rosenfeld, S. Dysinger, K. A. Lundgreen, L. Kuri-Cervantes, S. Adamski, A. Hicks, S. Korte, D. A. Oldridge, A. E. Baxter, J. R. Giles, M. E. Weirick, C. M. McAllister, J. Dougherty, S. Long, K. D’Andrea, J. T. Hamilton, M. R. Betts, E. T. Luning Prak, P. Bates, S. E. Hensley, A. R. Greenplate, and E. J. Wherry, Distinct antibody and memory B cell responses in SARS-CoV-2 naïve and recovered individuals following mRNA vaccination. Science immunology 6, (2021).

23. E. Piano Mortari, C. Russo, M. R. Vinci, S. Terreri, A. Fernandez Salinas, L. Piccioni, C. Alteri, L. Colagrossi, L. Coltella, S. Ranno, G. Linardos, M. Agosta, C. Albano, C. Agrati, C. Castilletti, S. Meschi, P. Romania, G. Roscilli, E. Pavoni, V. Camisa, A. Santoro, R. Brugaletta, N. Magnavita, A. Ruggiero, N. Cotugno, D. Amodio, M. L. Ciofi Degli Atti, D. Giorgio, N. Russo, G. Salvatori, T. Corsetti, F. Locatelli, C. F. Perno, S. Zaffina, and R. Carsetti, Highly Specific Memory B Cells Generation after the 2nd Dose of BNT162b2 Vaccine Compensate for the Decline of Serum Antibodies and Absence of Mucosal IgA. Cells 10, (2021).

24. G. E. Hartley, E. S. J. Edwards, P. M. Aui, N. Varese, S. Stojanovic, J. McMahon, A. Y. Peleg, I. Boo, H. E. Drummer, P. M. Hogarth, R. E. O’Hehir, and M. C. van Zelm, Rapid generation of durable B cell memory to SARS-CoV-2 spike and nucleocapsid proteins in COVID-19 and convalescence. Sci Immunol 5, (2020).

25. G. E. Hartley, E. S. J. Edwards, N. Varese, I. Boo, P. M. Aui, S. J. Bornheimer, P. M. Hogarth, H. E. Drummer, R. E. O’Hehir, and M. C. van Zelm, The second COVID-19 mRNA vaccine dose enhances the capacity of Spike-specific memory B cells to bind Omicron BA.2. Allergy, (2022).

26. J. M. Dan, J. Mateus, Y. Kato, K. M. Hastie, E. D. Yu, C. E. Faliti, A. Grifoni, S. I. Ramirez, S. Haupt, A. Frazier, C. Nakao, V. Rayaprolu, S. A. Rawlings, B. Peters, F. Krammer, V. Simon, E. O. Saphire, D. M. Smith, D. Weiskopf, A. Sette, and S. Crotty, Immunological memory to SARS-CoV-2 assessed for up to 8 months after infection. Science 371, (2021).

27. H. A. Fryer, G. E. Hartley, E. S. J. Edwards, N. Varese, I. Boo, S. J. Bornheimer, P. M. Hogarth, H. E. Drummer, R. E. O’Hehir, and M. C. van Zelm, COVID-19 Adenoviral Vector Vaccination Elicits a Robust Memory B Cell Response with the Capacity to Recognize Omicron BA.2 and BA.5 Variants. Journal of Clinical Immunology 43, 1506–1518 (2023).

28. N. H. Tan, R. S. G. Sablerolles, W. J. R. Rietdijk, A. Goorhuis, D. F. Postma, L. G. Visser, S. Bogers, D. Geers, L. M. Zaeck, M. P. G. Koopmans, V. A. S. H. Dalm, N. A. Kootstra, A. L. W. Huckriede, D. van Baarle, M. Lafeber, C. H. GeurtsvanKessel, R. D. de Vries, and P.-H. M. van der Kuy, Analyzing the immunogenicity of bivalent booster vaccinations in healthcare workers: The SWITCH ON trial protocol. Frontiers in Immunology 13, (2022).

29. R. R. Goel, M. Painter Mark, A. Apostolidis Sokratis, D. Mathew, W. Meng, M. Rosenfeld Aaron, A. Lundgreen Kendall, A. Reynaldi, S. Khoury David, A. Pattekar, S. Gouma, L. Kuri-Cervantes, P. Hicks, S. Dysinger, A. Hicks, H. Sharma, S. Herring, S. Korte, E. Baxter Amy, A. Oldridge Derek, R. Giles Josephine, E. Weirick Madison, M. McAllister Christopher, M. Awofolaju, N. Tanenbaum, M. Drapeau Elizabeth, J. Dougherty, S. Long, K. D’Andrea, T. Hamilton Jacob, M. McLaughlin, C. Williams Justine, S. Adamski, O. Kuthuru, n. null, I. Frank, R. Betts Michael, A. Vella Laura, A. Grifoni, D. Weiskopf, A. Sette, E. Hensley Scott, P. Davenport Miles, P. Bates, T. Luning Prak Eline, R. Greenplate Allison, and E. J. Wherry, mRNA vaccines induce durable immune memory to SARS-CoV-2 and variants of concern. Science 374, (2021).

30. N. H. Tan, D. Geers, R. S. G. Sablerolles, W. J. R. Rietdijk, A. Goorhuis, D. F. Postma, L. G. Visser, S. Bogers, L. L. A. van Dijk, L. Gommers, L. P. M. van Leeuwen, A. Boerma, S. H. Nijhof, K. A. van Dort, M. P. G. Koopmans, V. Dalm, M. Lafeber, N. A. Kootstra, A. L. W. Huckriede, D. van Baarle, L. M. Zaeck, C. H. GeurtsvanKessel, R. D. de Vries, and P. H. M. van der Kuy, Immunogenicity of bivalent omicron (BA.1) booster vaccination after different priming regimens in health-care workers in the Netherlands (SWITCH ON): results from the direct boost group of an open-label, multicentre, randomised controlled trial. Lancet Infect Dis 23, 901–913 (2023).

31. A. H. Ellebedy, K. J. Jackson, H. T. Kissick, H. I. Nakaya, C. W. Davis, K. M. Roskin, A. K. McElroy, C. M. Oshansky, R. Elbein, S. Thomas, G. M. Lyon, C. F. Spiropoulou, A. K. Mehta, P. G. Thomas, S. D. Boyd, and R. Ahmed, Defining antigen-specific plasmablast and memory B cell subsets in human blood after viral infection or vaccination. Nat Immunol 17, 1226–1234 (2016).

32. D. Lau, L. Y.-L. Lan, S. F. Andrews, C. Henry, K. T. Rojas, K. E. Neu, M. Huang, Y. Huang, B. DeKosky, A.-K. E. Palm, G. C. Ippolito, G. Georgiou, and P. C. Wilson, Low CD21 expression defines a population of recent germinal center graduates primed for plasma cell differentiation. Science immunology 2, (2017).

33. M. A. Berkowska, G. J. Driessen, V. Bikos, C. Grosserichter-Wagener, K. Stamatopoulos, A. Cerutti, B. He, K. Biermann, J. F. Lange, M. van der Burg, J. J. van Dongen, and M. C. van Zelm, Human memory B cells originate from three distinct germinal center-dependent and - independent maturation pathways. Blood 118, 2150–2158 (2011).

34. J. S. Buhre, T. Pongracz, I. Künsting, A. S. Lixenfeld, W. Wang, J. Nouta, S. Lehrian, F. Schmelter, H. B. Lunding, L. Dühring, C. Kern, J. Petry, E. L. Martin, B. Föh, M. Steinhaus, V. von Kopylow, C. Sina, T. Graf, J. Rahmöller, M. Wuhrer, and M. Ehlers, mRNA vaccines against SARS-CoV-2 induce comparably low long-term IgG Fc galactosylation and sialylation levels but increasing long-term IgG4 responses compared to an adenovirus-based vaccine. Frontiers in Immunology 13, (2023).

35. P. Irrgang, J. Gerling, K. Kocher, D. Lapuente, P. Steininger, K. Habenicht, M. Wytopil, S. Beileke, S. Schäfer, J. Zhong, G. Ssebyatika, T. Krey, V. Falcone, C. Schülein, A. S. Peter, K. Nganou-Makamdop, H. Hengel, J. Held, C. Bogdan, K. Überla, K. Schober, T. H. Winkler, and M. Tenbusch, Class switch toward noninflammatory, spike-specific IgG4 antibodies after repeated SARS-CoV-2 mRNA vaccination. Science Immunology 8, eade2798 (2023).

36. outbreak.info. 2023. BQ.1.1 Lineage Report, Vol. 2023. outbreak.info.

37. T. Tamura, J. Ito, K. Uriu, J. Zahradnik, I. Kida, Y. Anraku, H. Nasser, M. Shofa, Y. Oda, S. Lytras, N. Nao, Y. Itakura, S. Deguchi, R. Suzuki, L. Wang, M. S. T. M. Begum, S. Kita, H. Yajima, J. Sasaki, K. Sasaki-Tabata, R. Shimizu, M. Tsuda, Y. Kosugi, S. Fujita, L. Pan, D. Sauter, K. Yoshimatsu, S. Suzuki, H. Asakura, M. Nagashima, K. Sadamasu, K. Yoshimura, Y. Yamamoto, T. Nagamoto, G. Schreiber, K. Maenaka, H. Ito, N. Misawa, I. Kimura, M. Suganami, M. Chiba, R. Yoshimura, K. Yasuda, K. Iida, N. Ohsumi, A. P. Strange, O. Takahashi, K. Ichihara, Y. Shibatani, T. Nishiuchi, M. Kato, Z. Ferdous, H. Mouri, K. Shishido, H. Sawa, R. Hashimoto, Y. Watanabe, A. Sakamoto, N. Yasuhara, T. Suzuki, K. Kimura, Y. Nakajima, S. Nakagawa, J. Wu, K. Shirakawa, A. Takaori-Kondo, K. Nagata, Y. Kazuma, R. Nomura, Y. Horisawa, Y. Tashiro, Y. Kawai, T. Irie, R. Kawabata, C. Motozono, M. Toyoda, T. Ueno, T. Hashiguchi, T. Ikeda, T. Fukuhara, A. Saito, S. Tanaka, K. Matsuno, K. Takayama, K. Sato, and C. The Genotype to Phenotype Japan, Virological characteristics of the SARS-CoV-2 XBB variant derived from recombination of two Omicron subvariants. Nature Communications 14, 2800 (2023).

38. L. M. Zaeck, N. H. Tan, W. J. R. Rietdijk, D. Geers, R. S. G. Sablerolles, S. Bogers, L. L. A. v. Dijk, L. Gommers, L. P. M. v. Leeuwen, S. Rugebregt, A. Goorhuis, D. F. Postma, L. G. Visser, V. A. S. H. Dalm, M. Lafeber, N. A. Kootstra, A. L. W. Huckriede, B. L. Haagmans, D. v. Baarle, M. P. G. Koopmans, P. H. M. v. d. Kuy, C. H. GeurtsvanKessel, R. D. d. Vries, and S.-O. R. Group, Distinct COVID-19 vaccine combinations result in divergent immune responses. medRxiv, 2023.2008.2025.23294606 (2023).

39. M. E. Davis-Gardner, L. Lai, B. Wali, H. Samaha, D. Solis, M. Lee, A. Porter-Morrison, I. T. Hentenaar, F. Yamamoto, S. Godbole, Y. Liu, D. C. Douek, F. E.-H. Lee, N. Rouphael, A. Moreno, B. A. Pinsky, and M. S. Suthar, Neutralization against BA.2.75.2, BQ.1.1, and XBB from mRNA Bivalent Booster. New England Journal of Medicine 388, 183–185 (2022).

40. D. N. Springer, M. Bauer, I. Medits, J. V. Camp, S. W. Aberle, C. Burtscher, E. Höltl, L. Weseslindtner, K. Stiasny, and J. H. Aberle, Bivalent COVID-19 mRNA booster vaccination (BA.1 or BA.4/BA.5) increases neutralization of matched Omicron variants. npj Vaccines 8, 110 (2023).

41. G. E. Hartley, H. A. Fryer, P. A. Gill, I. Boo, S. J. Bornheimer, P. M. Hogarth, H. E. Drummer, R. E. O’Hehir, E. S. J. Edwards, and M. C. van Zelm, Third dose COVID-19 mRNA vaccine enhances IgG4 isotype switching and recognition of Omicron subvariants by memory B cells after mRNA but not adenovirus priming. bioRxiv, 2023.2009.2015.557929 (2023).

42. R. G. E. Krause, T. Moyo-Gwete, S. I. Richardson, Z. Makhado, N. P. Manamela, T. Hermanus, N. N. Mkhize, R. Keeton, N. Benede, M. Mennen, S. Skelem, F. Karim, K. Khan, C. Riou, N. A. B. Ntusi, A. Goga, G. Gray, W. Hanekom, N. Garrett, L.-G. Bekker, A. Groll, A. Sigal, P. L. Moore, W. A. Burgers, and A. Leslie, Infection pre-Ad26.COV2.S-vaccination primes greater class switching and reduced CXCR5 expression by SARS-CoV-2-specific memory B cells. npj Vaccines 8, 119 (2023).

43. M. E. Reincke, K. J. Payne, I. Harder, V. Strohmeier, R. E. Voll, K. Warnatz, and B. Keller, The Antigen Presenting Potential of CD21low B Cells. Frontiers in Immunology 11, (2020).

44. E. H. Gemma, A. F. Holly, A. G. Paul, B. Irene, J. B. Scott, P. M. Hogarth, E. D. Heidi, E. O. H. Robyn, S. J. E. Emily, and C. v. Z. Menno, Third dose COVID-19 mRNA vaccine enhances IgG4 isotype switching and recognition of Omicron subvariants by memory B cells after mRNA but not adenovirus priming. bioRxiv, 2023.2009.2015.557929 (2023).

45. M. Voysey, S. A. C. Clemens, S. A. Madhi, L. Y. Weckx, P. M. Folegatti, P. K. Aley, B. Angus, V. L. Baillie, S. L. Barnabas, Q. E. Bhorat, S. Bibi, C. Briner, P. Cicconi, E. A. Clutterbuck, A. M. Collins, C. L. Cutland, T. C. Darton, K. Dheda, C. Dold, C. J. A. Duncan, K. R. W. Emary, K. J. Ewer, A. Flaxman, L. Fairlie, S. N. Faust, S. Feng, D. M. Ferreira, A. Finn, E. Galiza, A. L. Goodman, C. M. Green, C. A. Green, M. Greenland, C. Hill, H. C. Hill, I. Hirsch, A. Izu, D. Jenkin, C. C. D. Joe, S. Kerridge, A. Koen, G. Kwatra, R. Lazarus, V. Libri, P. J. Lillie, N. G. Marchevsky, R. P. Marshall, A. V. A. Mendes, E. P. Milan, A. M. Minassian, A. McGregor, Y. F. Mujadidi, A. Nana, S. D. Padayachee, D. J. Phillips, A. Pittella, E. Plested, K. M. Pollock, M. N. Ramasamy, A. J. Ritchie, H. Robinson, A. V. Schwarzbold, A. Smith, R. Song, M. D. Snape, E. Sprinz, R. K. Sutherland, E. C. Thomson, M. E. Török, M. Toshner, D. P. J. Turner, J. Vekemans, T. L. Villafana, T. White, C. J. Williams, A. D. Douglas, A. V. S. Hill, T. Lambe, S. C. Gilbert, A. J. Pollard, M. Aban, K. W. M. Abeyskera, J. Aboagye, M. Adam, K. Adams, J. P. Adamson, G. Adewatan, S. Adlou, K. Ahmed, Y. Akhalwaya, S. Akhalwaya, A. Alcock, A. Ali, E. R. Allen, L. Allen, F. B. Alvernaz, F. S. Amorim, C. S. Andrade, F. Andritsou, R. Anslow, E. H. Arbe-Barnes, M. P. Ariaans, B. Arns, L. Arruda, L. Assad, P. D. A. Azi, L. D. A. Azi, G. Babbage, C. Bailey, K. F. Baker, M. Baker, N. Baker, P. Baker, I. Baleanu, D. Bandeira, A. Bara, M. A. S. Barbosa, D. Barker, G. D. Barlow, E. Barnes, A. S. Barr, J. R. Barrett, J. Barrett, K. Barrett, L. Bates, A. Batten, K. Beadon, E. Beales, R. Beckley, S. Belij-Rammerstorfer, J. Bell, D. Bellamy, S. Belton, A. Berg, L. Bermejo, E. Berrie, L. Berry, D. Berzenyi, A. Beveridge, K. R. Bewley, I. Bharaj, S. Bhikha, A. E. Bhorat, Z. E. Bhorat, E. M. Bijker, S. Birch, G. Birch, K. Birchall, A. Bird, O. Bird, K. Bisnauthsing, M. Bittaye, L. Blackwell, R. Blacow, H. Bletchly, C. L. Blundell, S. R. Blundell, P. Bodalia, E. Bolam, E. Boland, D. Bormans, N. Borthwick, F. Bowring, A. Boyd, P. Bradley, T. Brenner, A. Bridges-Webb, P. Brown, C. Brown, C. Brown-O’Sullivan, S. Bruce, E. Brunt, W. Budd, Y. A. Bulbulia, M. Bull, J. Burbage, A. Burn, K. R. Buttigieg, N. Byard, I. Cabrera Puig, A. Calvert, S. Camara, M. Cao, F. Cappuccini, R. Cardona, J. R. Cardoso, M. Carr, M. W. Carroll, A. Carson-Stevens, Y. d. M. Carvalho, H. R. Casey, P. Cashen, T. R. Y. Castro, L. C. Castro, K. Cathie, A. Cavey, J. Cerbino-Neto, L. F. F. Cezar, J. Chadwick, C. Chanice, D. Chapman, S. Charlton, K. S. Cheliotis, I. Chelysheva, O. Chester, E. Chiplin, S. Chita, J.-S. Cho, L. Cifuentes, E. Clark, M. Clark, R. Colin-Jones, S. L. K. Collins, H. Colton, C. P. Conlon, S. Connarty, N. Coombes, C. Cooper, R. Cooper, L. Cornelissen, T. Corrah, C. A. Cosgrove, F. B. Costa, T. Cox, W. E. M. Crocker, S. Crosbie, D. Cullen, D. R. M. F. Cunha, C. J. Cunningham, F. C. Cuthbertson, D. M. da Costa, S. N. F. Da Guarda, L. P. da Silva, A. C. da Silva Moraes, B. E. Damratoski, Z. Danos, M. T. D. C. Dantas, M. S. Datoo, C. Datta, M. Davids, S. L. Davies, K. Davies, H. Davies, S. Davies, J. Davies, E. J. Davis, J. Davis, J. A. M. de Carvalho, J. De Jager, S. de Jesus Jnr, L. M. De Oliveira Kalid, D. Dearlove, T. Demissie, A. Desai, S. Di Marco, C. Di Maso, T. Dinesh, C. Docksey, T. Dong, F. R. Donnellan, T. G. Dos Santos, T. G. Dos Santos, E. P. Dos Santos, N. Douglas, C. Downing, J. Drake, R. Drake-Brockman, R. Drury, J. Du Plessis, S. J. Dunachie, A. Duncan, N. J. W. Easom, M. Edwards, N. J. Edwards, F. Edwards, O. M. El Muhanna, S. C. Elias, B. Ellison-Handley, M. J. Elmore, M. R. English, A. Esmail, Y. M. Essack, M. Farooq, S. Fedosyuk, S. Felle, S. Ferguson, C. Ferreira Da Silva, S. Field, R. Fisher, J. Fletcher, H. Fofie, H. Fok, K. J. Ford, R. Fothergill, J. Fowler, P. H. A. Fraiman, E. Francis, M. M. Franco, J. Frater, M. S. M. Freire, S. H. Fry, S. Fudge, R. Furlan Filho, J. Furze, M. Fuskova, P. Galian-Rubio, H. Garlant, M. Gavrila, K. A. Gibbons, C. Gilbride, H. Gill, K. Godwin, K. Gokani, M. L. F. Gonçalves, I. G. S. Gonzalez, J. Goodall, J. Goodwin, A. Goondiwala, K. Gordon-Quayle, G. Gorini, A. Goyanna, J. Grab, L. Gracie, J. Green, N. Greenwood, J. Greffrath, M. M. Groenewald, A. Gunawardene, G. Gupta, M. Hackett, B. Hallis, M. Hamaluba, E. Hamilton, J. Hamlyn, D. Hammersley, A. T. Hanrath, B. Hanumunthadu, S. A. Harris, C. Harris, T. D. Harrison, D. Harrison, T. A. Harris-Wright, T. C. Hart, B. Hartnell, J. Haughney, S. Hawkins, L. Y. M. Hayano, I. Head, P. T. Heath, J. A. Henry, M. Hermosin Herrera, D. B. Hettle, C. Higa, J. Hill, G. Hodges, S. Hodgson, E. Horne, M. M. Hou, C. F. Houlihan, E. Howe, N. Howell, J. Humphreys, H. E. Humphries, K. Hurley, C. Huson, C. Hyams, A. Hyder-Wright, S. Ikram, A. Ishwarbhai, P. Iveson, V. Iyer, F. Jackson, S. Jackson, S. Jaumdally, H. Jeffers, N. Jesudason, C. Jones, C. Jones, K. Jones, E. Jones, M. R. Jorge, A. Joshi, E. A. M. S. Júnior, R. Kailath, F. Kana, A. Kar, K. Karampatsas, M. Kasanyinga, L. Kay, J. Keen, J. Kellett Wright, E. J. Kelly, D. Kelly, D. M. Kelly, S. Kelly, D. Kerr, L. Khan, B. Khozoee, A. Khurana, S. Kidd, A. Killen, J. Kinch, P. Kinch, L. D. W. King, T. B. King, L. Kingham, P. Klenerman, D. M. Kluczna, F. Knapper, J. C. Knight, D. Knott, S. Koleva, P. M. Lages, M. Lang, G. Lang, C. W. Larkworthy, J. P. J. Larwood, R. Law, A. M. Lawrie, E. M. Lazarus, A. Leach, E. A. Lees, A. Lelliott, N.-M. Lemm, A. E. R. Lessa, S. Leung, Y. Li, A. M. Lias, K. Liatsikos, A. Linder, S. Lipworth, S. Liu, X. Liu, A. Lloyd, S. Lloyd, L. Loew, R. Lopez Ramon, L. B. Lora, K. G. Luz, J. C. MacDonald, G. MacGregor, M. Madhavan, D. O. Mainwaring, E. Makambwa, R. Makinson, M. Malahleha, R. Malamatsho, G. Mallett, N. Manning, K. Mansatta, T. Maoko, S. Marinou, E. Marlow, G. N. Marques, P. Marriott, R. P. Marshall, J. L. Marshall, M. Masenya, M. Masilela, S. K. Masters, M. Mathew, H. Matlebjane, K. Matshidiso, O. Mazur, A. Mazzella, H. McCaughan, J. McEwan, J. McGlashan, L. McInroy, N. McRobert, S. McSwiggan, C. Megson, S. Mehdipour, W. Meijs, R. N. Õ. Mendonça, A. J. Mentzer, A. C. F. Mesquita, P. Miralhes, N. Mirtorabi, C. Mitton, S. Mnyakeni, F. Moghaddas, K. Molapo, M. Moloi, M. Moore, M. Moran, E. Morey, R. Morgans, S. J. Morris, S. Morris, H. Morrison, F. Morselli, G. Morshead, R. Morter, L. Mottay, A. Moultrie, N. Moyo, M. Mpelembue, S. Msomi, Y. Mugodi, E. Mukhopadhyay, J. Muller, A. Munro, S. Murphy, P. Mweu, C. Myerscough, G. Naik, K. Naker, E. Nastouli, B. Ndlovu, E. Nikolaou, C. Njenga, H. C. Noal, A. Noé, G. Novaes, F. L. Nugent, G. L. A. Nunes, K. O’Brien, D. O’Connor, S. Oelofse, B. Oguti, V. Olchawski, N. J. Oldfield, M. G. Oliveira, C. Oliveira, I. S. Q. Oliveira, A. Oommen-Jose, A. Oosthuizen, P. O’Reilly, P. J. O’Reilly, P. Osborne, D. R. J. Owen, L. Owen, D. Owens, N. Owino, M. Pacurar, B. V. B. Paiva, E. M. F. Palhares, S. Palmer, H. M. R. T. Parracho, K. Parsons, D. Patel, B. Patel, F. Patel, M. Patrick-Smith, R. O. Payne, Y. Peng, E. J. Penn, A. Pennington, M. P. Peralta Alvarez, B. P. Pereira Stuchi, A. L. Perez, T. Perinpanathan, J. Perring, R. Perumal, S. Y. Petkar, T. Philip, J. Phillips, M. K. Phohu, L. Pickup, S. Pieterse, J. M. Pinheiro, J. Piper, D. Pipini, M. Plank, S. Plant, S. Pollard, J. Pooley, A. Pooran, I. Poulton, C. Powers, F. B. Presa, D. A. Price, V. Price, M. R. Primeira, P. C. Proud, S. Provstgaard-Morys, S. Pueschel, D. Pulido, S. Quaid, R. Rabara, K. Radia, D. Rajapaska, T. Rajeswaran, L. Ramos, A. S. F. Ramos, F. Ramos Lopez, T. Rampling, J. Rand, H. Ratcliffe, T. Rawlinson, D. Rea, B. Rees, M. Resuello-Dauti, E. Reyes Pabon, S. Rhead, T. Riaz, M. Ricamara, A. Richards, A. Richter, N. Ritchie, A. J. Ritchie, A. J. Robbins, H. Roberts, R. E. Robinson, S. Roche, C. Rollier, L. Rose, A. L. Ross Russell, L. Rossouw, S. Royal, I. Rudiansyah, K. Ryalls, C. Sabine, S. Saich, J. C. Sale, A. M. Salman, N. Salvador, S. Salvador, M. D. Sampaio, A. D. Samson, A. Sanchez-Gonzalez, H. Sanders, K. Sanders, E. Santos, M. F. S. Santos Guerra, I. Satti, J. E. Saunders, C. Saunders, A. B. A. Sayed, I. Schim van der Loeff, A. B. Schmid, E. Schofield, G. R. Screaton, S. Seddiqi, R. R. Segireddy, R. Senger, S. Serrano, I. Shaik, H. R. Sharpe, K. Sharrocks, R. Shaw, A. Shea, E. Sheehan, A. Shepherd, F. Shiham, S. E. Silk, L. Silva-Reyes, L. B. T. D. Silveira, M. B. V. Silveira, N. Singh, J. Sinha, D. T. Skelly, D. C. Smith, N. Smith, H. E. Smith, D. J. Smith, C. C. Smith, A. S. Soares, C. Solórzano, G. L. Sorio, K. Sorley, T. Sosa-Rodriguez, C. M. C. D. L. Souza, B. S. D. F. Souza, A. R. Souza, T. Souza Lopez, L. Sowole, A. J. Spencer, L. Spoors, L. Stafford, I. Stamford, R. Stein, L. Stockdale, L. V. Stockwell, L. H. Strickland, A. Stuart, A. Sturdy, N. Sutton, A. Szigeti, A. Tahiri-Alaoui, R. Tanner, C. Taoushanis, A. W. Tarr, R. Tarrant, K. Taylor, U. Taylor, I. J. Taylor, J. Taylor, R. te Water Naude, K. Templeton, Y. Themistocleous, A. Themistocleous, M. Thomas, K. Thomas, T. M. Thomas, A. Thombrayil, J. Thompson, F. Thompson, A. Thompson, A. Thompson, K. Thompson, V. Thornton-Jones, L. H. S. Thotusi, P. J. Tighe, L. A. Tinoco, G. F. Tiongson, B. Tladinyane, M. Tomasicchio, A. Tomic, S. Tonks, J. Towner, N. Tran, J. A. Tree, G. Trillana, C. Trinham, R. Trivett, A. Truby, B. L. Tsheko, P. Tubb, A. Turabi, R. Turner, C. Turner, N. Turner, B. Tyagi, M. Ulaszewska, B. R. Underwood, S. van Eck, R. Varughese, D. Verbart, M. K. Verheul, I. Vichos, T. A. Vieira, G. Walker, L. Walker, M. E. Wand, T. Wardell, G. M. Warimwe, S. C. Warren, B. Watkins, M. E. E. Watson, E. Watson, S. Webb, A. Webster, J. Welch, Z. Wellbelove, J. H. Wells, A. J. West, B. White, C. White, R. White, P. Williams, R. L. Williams, S. Willingham, R. Winslow, D. Woods, M. Woodyer, A. T. Worth, D. Wright, M. Wroblewska, A. Yao, Y. T. N. Yim, M. B. Zambrano, R. L. Zimmer, D. Zizi, and P. Zuidewind, Single-dose administration and the influence of the timing of the booster dose on immunogenicity and efficacy of ChAdOx1 nCoV-19 (AZD1222) vaccine: a pooled analysis of four randomised trials. The Lancet 397, 881–891 (2021).

46. World Health Organisation. 2022. Interim recommendations for the use of the Janssen Ad26.COV2.S (COVID-19) vaccine. In COVID-19: Vaccines, Vol. 2024.

47. F. X. Heinz, and K. Stiasny, Distinguishing features of current COVID-19 vaccines: knowns and unknowns of antigen presentation and modes of action. npj Vaccines 6, 104 (2021).

48. J. E. Bowen, Y.-J. Park, C. Stewart, J. T. Brown, W. K. Sharkey, A. C. Walls, A. Joshi, K. R. Sprouse, M. McCallum, M. A. Tortorici, N. M. Franko, J. K. Logue, I. G. Mazzitelli, A. W. Nguyen, R. P. Silva, Y. Huang, J. S. Low, J. Jerak, S. W. Tiles, K. Ahmed, A. Shariq, J. M. Dan, Z. Zhang, D. Weiskopf, A. Sette, G. Snell, C. M. Posavad, N. T. Iqbal, J. Geffner, A. Bandera, A. Gori, F. Sallusto, J. A. Maynard, S. Crotty, W. C. Van Voorhis, C. Simmerling, R. Grifantini, H. Y. Chu, D. Corti, and D. Veesler, SARS-CoV-2 spike conformation determines plasma neutralizing activity elicited by a wide panel of human vaccines. Science Immunology 7, eadf1421 (2022).

49. R. K. Suryawanshi, T. Y. Taha, M. McCavitt-Malvido, I. Silva, M. M. Khalid, A. M. Syed, I. P. Chen, P. Saldhi, B. Sreekumar, M. Montano, K. Foresythe, T. Tabata, G. R. Kumar, A. Sotomayor-Gonzalez, V. Servellita, A. Gliwa, J. Nguyen, N. Kojima, T. Arellanor, A. Bussanich, V. Hess, M. Shacreaw, L. Lopez, M. Brobeck, F. Turner, Y. Wang, S. Ghazarian, G. Davis, D. Rodriguez, J. Doudna, L. Spraggon, C. Y. Chiu, and M. Ott, Previous exposure to Spike-providing parental strains confers neutralizing immunity to XBB lineage and other SARS-CoV-2 recombinants in the context of vaccination. Emerging Microbes & Infections 12, 2270071 (2023).

50. W. E. Purtha, T. F. Tedder, S. Johnson, D. Bhattacharya, and M. S. Diamond, Memory B cells, but not long-lived plasma cells, possess antigen specificities for viral escape mutants. Journal of Experimental Medicine 208, 2599–2606 (2011).

51. A. Sokal, G. Barba-Spaeth, L. Hunault, I. Fernández, M. Broketa, A. Meola, S. Fourati, I. Azzaoui, A. Vandenberghe, P. Lagouge-Roussey, M. Broutin, A. Roeser, M. Bouvier-Alias, E. Crickx, L. Languille, M. Fournier, M. Michel, B. Godeau, S. Gallien, G. Melica, Y. Nguyen, F. Canoui-Poitrine, F. Pirenne, J. Megret, J.-M. Pawlotsky, S. Fillatreau, C.-A. Reynaud, J.-C. Weill, F. A. Rey, P. Bruhns, M. Mahévas, and P. Chappert, SARS-CoV-2 Omicron BA.1 breakthrough infection drives late remodeling of the memory B cell repertoire in vaccinated individuals. Immunity 56, 2137–2151.e2137 (2023).

52. K. Röltgen, S. C. A. Nielsen, O. Silva, S. F. Younes, M. Zaslavsky, C. Costales, F. Yang, O. F. Wirz, D. Solis, R. A. Hoh, A. Wang, P. S. Arunachalam, D. Colburg, S. Zhao, E. Haraguchi, A. S. Lee, M. M. Shah, M. Manohar, I. Chang, F. Gao, V. Mallajosyula, C. Li, J. Liu, M. J. Shoura, S. B. Sindher, E. Parsons, N. J. Dashdorj, N. D. Dashdorj, R. Monroe, G. E. Serrano, T. G. Beach, R. S. Chinthrajah, G. W. Charville, J. L. Wilbur, J. N. Wohlstadter, M. M. Davis, B. Pulendran, M. L. Troxell, G. B. Sigal, Y. Natkunam, B. A. Pinsky, K. C. Nadeau, and S. D. Boyd, Immune imprinting, breadth of variant recognition, and germinal center response in human SARS-CoV-2 infection and vaccination. Cell 185, 1025–1040.e1014 (2022).

53. J. S. Turner, J. A. O’Halloran, E. Kalaidina, W. Kim, A. J. Schmitz, J. Q. Zhou, T. Lei, M. Thapa, R. E. Chen, J. B. Case, F. Amanat, A. M. Rauseo, A. Haile, X. Xie, M. K. Klebert, T. Suessen, W. D. Middleton, P.-Y. Shi, F. Krammer, S. A. Teefey, M. S. Diamond, R. M. Presti, and A. H. Ellebedy, SARS-CoV-2 mRNA vaccines induce persistent human germinal centre responses. Nature 596, 109–113 (2021).

54. C. C. Traut, and J. N. Blankson, Bivalent mRNA vaccine-elicited SARS-CoV-2 specific T cells recognise the omicron XBB sublineage. The Lancet Microbe 4, e388 (2023).

55. N. Baumgarth, How specific is too specific? B-cell responses to viral infections reveal the importance of breadth over depth. Immunological Reviews 255, 82–94 (2013).

56. N. Patel, J. F. Trost, M. Guebre-Xabier, H. Zhou, J. Norton, D. Jiang, Z. Cai, M. Zhu, A. M. Marchese, A. M. Greene, R. M. Mallory, R. Kalkeri, F. Dubovsky, and G. Smith, XBB.1.5 spike protein COVID-19 vaccine induces broadly neutralizing and cellular immune responses against EG.5.1 and emerging XBB variants. Scientific Reports 13, 19176 (2023).

57. M. A. Tortorici, A. Addetia, A. J. Seo, J. Brown, K. R. Sprouse, J. Logue, E. Clark, N. Franko, H. Chu, and D. Veesler, Persistent immune imprinting after XBB.1.5 COVID vaccination in humans. bioRxiv, 2023.2011.2028.569129 (2023).

58. R. S. G. Sablerolles, W. J. R. Rietdijk, A. Goorhuis, D. F. Postma, L. G. Visser, D. Geers, K. S. Schmitz, H. M. Garcia Garrido, M. P. G. Koopmans, V. A. S. H. Dalm, N. A. Kootstra, A. L. W. Huckriede, M. Lafeber, D. van Baarle, C. H. GeurtsvanKessel, R. D. de Vries, and P. H. M. van der Kuy, Immunogenicity and Reactogenicity of Vaccine Boosters after Ad26.COV2.S Priming. New England Journal of Medicine 386, 951–963 (2022).

59. D. Geers, M. C. Shamier, S. Bogers, G. den Hartog, L. Gommers, N. N. Nieuwkoop, K. S. Schmitz, L. C. Rijsbergen, J. A. T. van Osch, E. Dijkhuizen, G. Smits, A. Comvalius, D. van Mourik, T. G. Caniels, M. J. van Gils, R. W. Sanders, B. B. Oude Munnink, R. Molenkamp, H. J. de Jager, B. L. Haagmans, R. L. de Swart, M. P. G. Koopmans, R. S. van Binnendijk, R. D. de Vries, and C. H. GeurtsvanKessel, SARS-CoV-2 variants of concern partially escape humoral but not T cell responses in COVID-19 convalescent donors and vaccine recipients. Science Immunology 6, eabj1750 (2021).

60. E. S. J. Edwards, J. J. Bosco, P. M. Aui, R. G. Stirling, P. U. Cameron, J. Chatelier, F. Hore-Lacy, R. E. O’Hehir, and M. C. van Zelm, Predominantly Antibody-Deficient Patients With Non-infectious Complications Have Reduced Naive B, Treg, Th17, and Tfh17 Cells. Frontiers in Immunology 10, (2019).

