## Supplementary Material for "Bivalent COVID-19 vaccines boost the capacity of pre-existing SARS-CoV-2-specific memory B cells to cross-recognize Omicron subvariants"

**SUPPLEMENTARY MATERIALS:**

**Supplementary Tables (n=3) and Supplementary Figures (n=3)**

**Supplementary Table 1. RBD-specific Bmem responses to 4th dose COVID-19 booster**

| | **Bmem numbers** (Fold change post-dose 4) | | |
|---|---|---|---|
| **Booster type** | **WH1-specific** | **BA.1-specific** | **BA.5-specific** |
| **Monovalent (n=18)** | 1.35 | 1.50 | Not measured |
| **BA.1 bivalent (n=33)** | 1.48 | 1.38 | Not measured |
| **BA.5 bivalent (n=21)** | 1.46 | Not measured | 1.42 |

### Supplementary Table 2. Antibody panel compositions

| | **Fluorochrome** | | | | | | | | | | | | | | | | | | | |
|---|---|---|---|---|---|---|---|---|---|---|---|---|---|---|---|---|---|---|---|---|
| **Tube** | BUV 395 | BUV 496 | BUV 615 | BUV 737 | BUV 805 | BV421 | cFluor V450 | BV 480 | BV605 | BV650 | BV711 | BV786 | FITC | BB700/ PerCP Cy-5.5 | PE | PE-Vio615 | cFluor BYG710 | PE-Cy7/ BYG781 | APC | APC-Cy7/ ViaDye Red |
| **1. Trucount** | — | — | — | — | — | — | — | — | — | — | — | — | CD3 | CD45 | CD16 + CD56 | — | — | CD4 | CD19 | CD8a |
| **2a. Bmem (mono- and BA.1 bivalent recipients)** | WH1 RBD | BA.5 RBD | — | BA.1 RBD | CD3 | WH1 RBD | IgM | BA.1 RBD | CD38 | BQ.1.1 RBD | CD21 | CD71 | IgG2 + IgG3 | IgD | IgG1 + IgG2 | IgA | CD19 | CD27 | IgG4 | Fixable viability |
| **2b. Bmem (BA.5 bivalent recipients)** | WH1 RBD | BA.1 RBD | XBB.1.5 RBD | BA.5 RBD | CD3 | WH1 RBD | IgM | BA.5 RBD | CD38 | BQ.1.1 RBD | CD21 | CD71 | IgG2 + IgG3 | IgD | IgG1 + IgG2 | IgA | CD19 | CD27 | IgG4 | Fixable viability |
| **3a. Strep control (mono- and BA.1 bivalent recipients)** | Strep | Strep | — | Strep | CD3 | Strep | — | Strep | — | Strep | — | — | — | IgD | — | — | CD19 | CD27 | — | Fixable viability |
| **3b. Strep control (BA.5 bivalent recipients)** | Strep | Strep | Strep | Strep | CD3 | Strep | — | Strep | — | Strep | — | — | — | IgD | — | — | CD19 | CD27 | — | Fixable viability |

**Supplementary Table 3. Antibody details**

| Marker | Fluoro-chrome | Clone | Vendor | Cat. number | Volume (µL)/ 100µL test | Tube |
|---|---|---|---|---|---|---|
| CD3 | BUV805 | UCHT1 | BD Biosciences | 612896 | 2.5 | 2, 3 |
| CD3 | FITC | SK7 | BD Biosciences | 662995* | 46ng | 1 |
| CD4 | PE-Cy7 | SK3 | BD Biosciences | 662995* | 30ng | 1 |
| CD8a | APC-Cy7 | SK1 | BD Biosciences | 662995* | 126ng | 1 |
| CD16 | PE | B73.1 | BD Biosciences | 662995* | 33ng | 1 |
| CD19 | APC | SJ25C1 | BD Biosciences | 662995* | 46ng | 1 |
| CD19 | cFluor BYG710 | HIB19 | Cytek Biosciences | SKU-R7-20010 | 1 | 2, 3 |
| CD21 | BV711 | B-ly4 | BD Biosciences | 563163 | 5 | 2 |
| CD27 | cFluor BYG781 | O323 | Cytek Biosciences | Custom | 0.5 | 2, 3 |
| CD38 | BV605 | HB7 | BD Biosciences | 562665 | 0.2 | 2 |
| CD45 | PerCP Cy-5.5 | 2D1 | BD Biosciences | 662995* | 120ng | 1 |
| CD56 | PE | NCAM16.2 | BD Biosciences | 662995* | 22ng | 1 |
| CD71 | BV786 | M-A712 | BD Biosciences | 563768 | 1 | 2 |
| Fixable Viability | ViaDye Red | - | Cytek Biosciences | SKU-R7-60008 | 0.025 | 2, 3 |
| IgA | PE-Vio615 | REA1014 | Miltenyi Biotec | 130-116-882 | 1.5 | 2 |
| IgD | BB700 | IA6-2 | BD Biosciences | 566538 | 1 | 2, 3 |
| IgG1 | PE | G17-1 | BD Biosciences | 624049 | 0.1 | 2 |
| IgG2 | FITC | HP6002 | BD Biosciences | 624045 | 0.5 | 2 |
| IgG2 | PE | HP6002 | BD Biosciences | 624049 | 1 | 2 |
| IgG3 | FITC | HP6047 | BD Biosciences | 624045 | 0.5 | 2 |
| IgG4 | APC | SAG4 | Cytognos | CYT-IGG4AP | 2 | 2 |
| IgM | cFluor V450 | MHM88 | Cytek Biosciences | Custom | 0.25 | 2 |
| Streptavidin | BUV395 | - | BD Biosciences | 564176 | 0.67 | 2, 3 |
| Streptavidin | BUV496 | - | BD Biosciences | 612961 | 0.67 | 2, 3 |
| Streptavidin | BUV615 | - | BD Biosciences | 613013 | 0.67 | 2b, 3b only |
| Streptavidin | BUV737 | - | BD Biosciences | 612775 | 0.67 | 2, 3 |
| Streptavidin | BV421 | - | BioLegend | 405225 | 0.13 | 2, 3 |
| Streptavidin | BV480 | - | BD Biosciences | 564876 | 0.67 | 2, 3 |
| Streptavidin | BV650 | - | BioLegend | 563855 | 0.13 | 2, 3 |
| *Antibodies part of the Multitest$^{TM}$ 6-color TBNK kit (BD Biosciences, Cat. number 662967) | | | | | | |

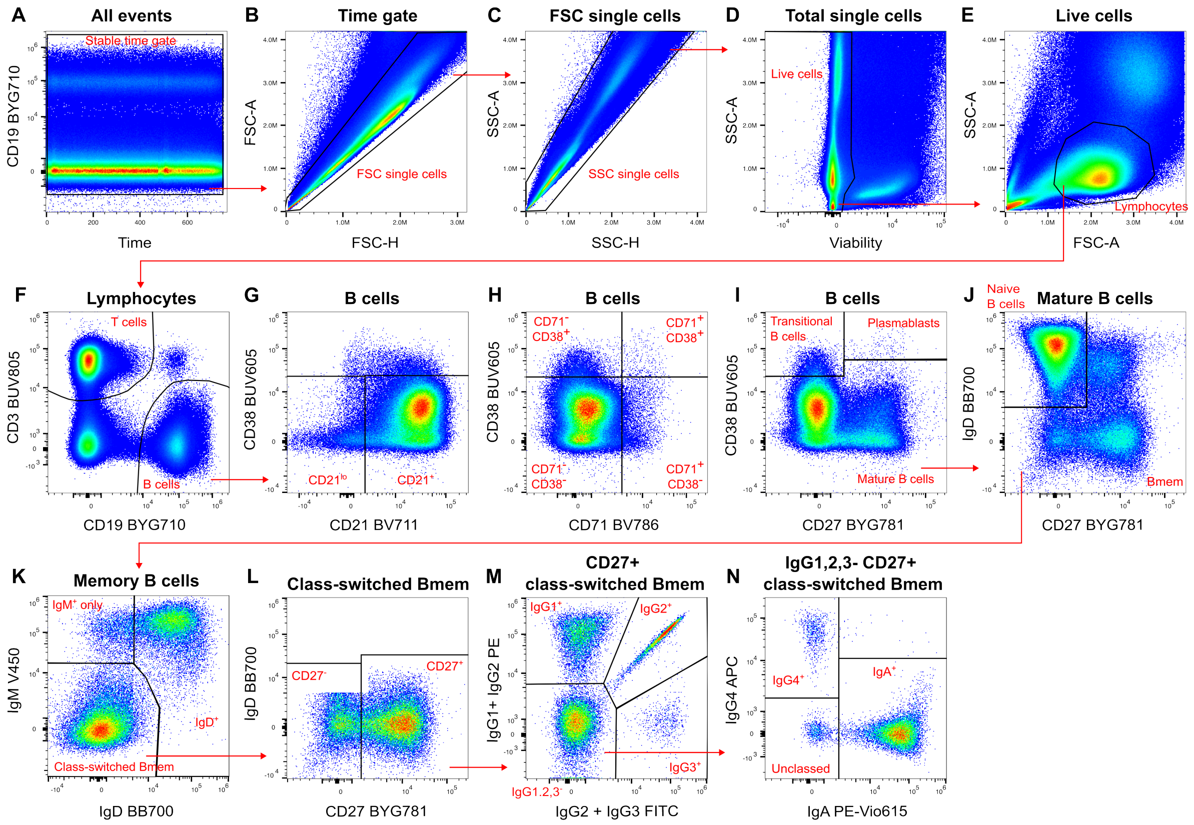


**Supplementary Figure 1. B-cell gating strategy.** (**A**) All events were gated on CD19 vs time to ensure a steady signal across acquisition time and to exclude aggregates. (**B-C**) Doublets were excluded on FSC-A vs FSC-H, and subsequently SSC-A vs SSC-H. (**D**) Dead cells were excluded on SSC-A vs viability dye. (**E**) Lymphocytes were gated as $SSC^{lo}FSC^{mid}$. (**F**) B cells were gated as $CD19^{+}CD3^{-}$. (**G**) $CD21^{lo}$ and $CD21^{+}$ B cells were gated vs CD38. (**H**) CD38 and CD71 expression on B cells was gated in quadrants. (**I**) Mature and transitional B cells and plasmablasts were gated on CD38 vs CD27. (**J**) Memory B cells (Bmem) and naive B cells were gated on IgD vs CD27 within mature B cells. (**K**) Class-switched, $IgD^{+}$, and $IgM^{+}$ only mature Bmem were gated on IgM vs IgD. (**L**) $CD27^{-}$ and $CD27^{+}$ class-switched mature Bmem were gated vs IgD. (**M**) $IgG1^{+}$, $IgG2^{+}$, $IgG3^{+}$, and $IgG1,2,3^{-}$ class-switched mature Bmem were gated within $CD27^{+}$ cells (and $CD27^{-}$, plot not shown). (**N**) $IgG4^{+}$, $IgA^{+}$, and unclassed mature Bmem were gated within $IgG1,2,3^{-}CD27^{+}$ cells (and $CD27^{-}$, plot not shown). Representative plots from BA.1 bivalent booster donor, post-dose 4.

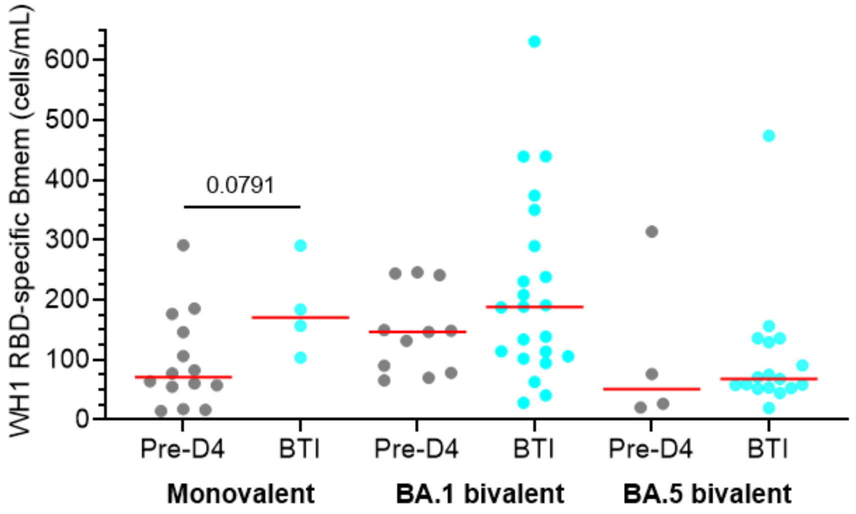


**Supplementary Figure 2. The effect of breakthrough infections on the RBD-specific Bmem response.** Absolute numbers of WH1 RBD-specific Bmem pre-dose 4, divided by booster type group and breakthrough infection (BTI) status. Blue dots denote any confirmed SARS-CoV-2 BTI before pre-dose 4 sampling. Monovalent, n=18; BA.1 bivalent, n=33; BA.5 bivalent, n=21. Solid lines depict median values. Mann-Whitney test for unpaired data.

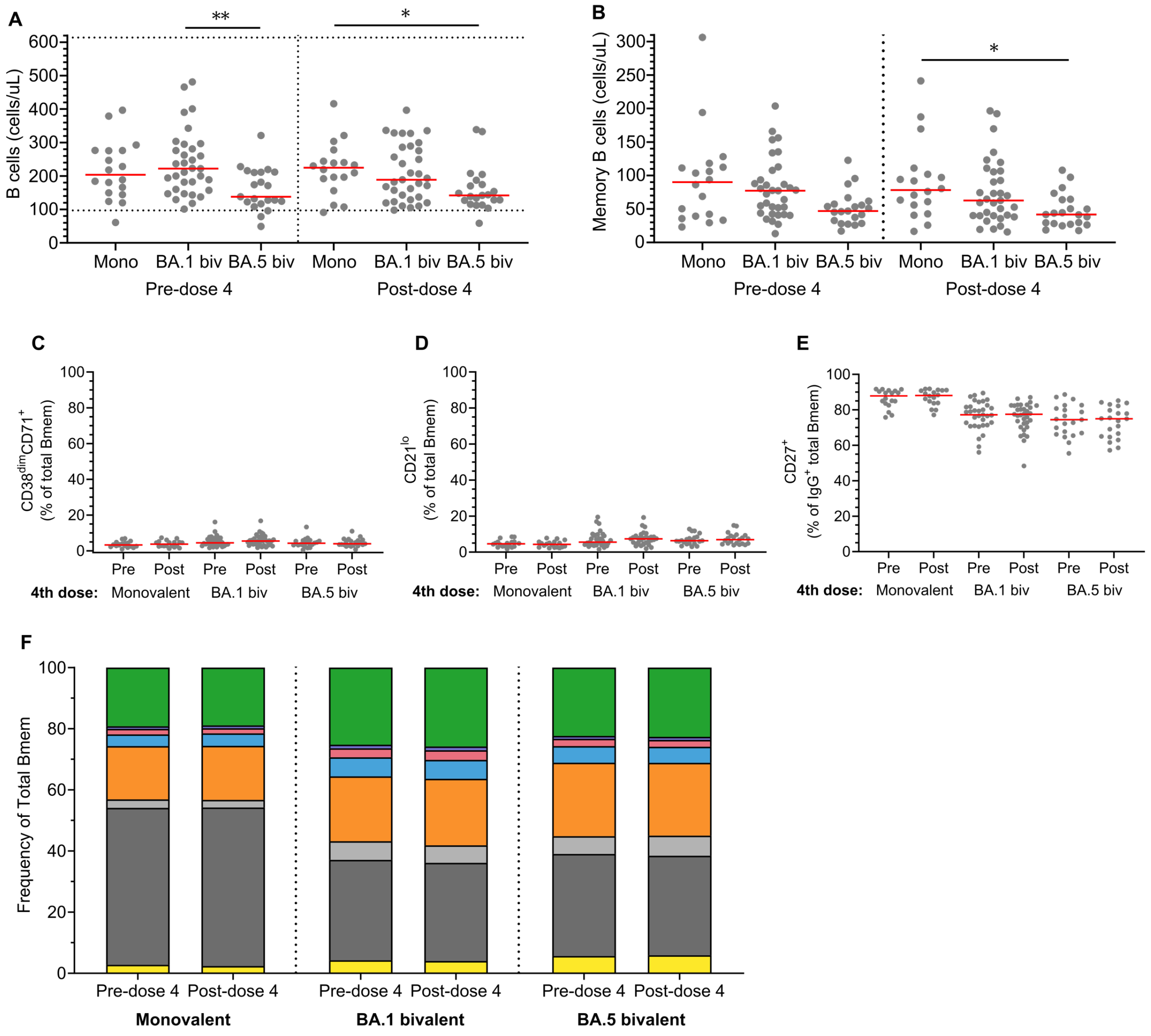


**Supplementary Figure 3. Total memory B cell numbers and Ig isotypes before and after monovalent or bivalent dose 4.** (**A**) Absolute numbers of total B cells and (**B**) memory B cells (Bmem) before and after a monovalent, BA.1 bivalent, or BA.5 bivalent dose 4. Dotted lines in (**A**) denote $5^{th}$ and $95^{th}$ percentiles of B-cell numbers from healthy controls previously published (60). (**C**) Frequencies of $CD38^{dim}CD71^{+}$, (**D**) $CD21^{lo}CD38^{dim}$, and (**E**) $CD27^{+}IgG^{+}$ Bmem pre- and 4-weeks post-monovalent, BA.1 bivalent, or BA.5 bivalent dose 4. (**F**) Frequencies of $IgG1^{+}$, $IgG2^{+}$, $IgG3^{+}$, $IgG4^{+}$, $IgA^{+}$, $IgD^{+}$, and $IgM^{+}$-only within total Bmem pre- and 4-weeks post-monovalent, BA.1 bivalent, or BA.5 bivalent dose 4. Monovalent, n=18; BA.1 bivalent, n=33; BA.5 bivalent, n=21. Solid lines depict median values. Kruskal-Wallis test with Dunn's multiple comparisons for multiple unpaired groups, and Wilcoxon matched-pairs signed rank test for paired data. Only significant differences shown. *$p<0.05$, **$p<0.01$.
